# CBFA2T3-GLIS2-dependent pediatric acute megakaryoblastic leukemia is driven by GLIS2 and sensitive to Navitoclax

**DOI:** 10.1101/2022.12.16.520129

**Authors:** Mathieu Neault, Charles-Étienne Lebert-Ghali, Marilaine Fournier, Caroline Capdevielle, Elizabeth A.R. Garfinkle, Alyssa Obermayer, Anitria Cotton, Karine Boulay, Christina Sawchyn, Kamy H. Nguyen, Béatrice Assaf, François E. Mercier, Jean-Sébastien Delisle, Elliot A. Drobetsky, Laura Hulea, Timothy I. Shaw, Johannes Zuber, Tanja A. Gruber, Heather J. Melichar, Frédérick A. Mallette

## Abstract

Pediatric acute megakaryoblastic leukemia (AMKL) is an aggressive, uncurable blood cancer associated with poor therapeutic response and high mortality. We developed CBFA2T3-GLIS2-driven mouse models of AMKL that recapitulate the phenotypic and transcriptional signatures of the human disease. We show that an activating Ras mutation, which occurs in human AMKL, increased the penetrance and decreased the latency of CBF2AT3-GLIS2-driven AMKL. CBFA2T3-GLIS2 and GLIS2 modulate similar transcriptional networks. We uncover the dominant oncogenic properties of GLIS2, which trigger AMKL in cooperation with oncogenic Ras. We find that both CBFA2T3-GLIS2 and GLIS2 alter the expression of numerous BH3-only proteins, causing AMKL cell sensitivity to the BCL-2 inhibitor navitoclax both *in vitro* and *in vivo*, suggesting a novel therapeutic option for pediatric patients suffering from CBFA2T3-GLIS2-driven AMKL.

**Key points:** GLIS2 cooperates with activated Nras to promote the development of acute megakaryoblastic leukemia.

CBFA2T3-GLIS2 and GLIS2 alter the expression of BCL2 family members rendering AMKL cells sensitive to navitoclax.

## Introduction

Pediatric acute megakaryoblastic leukemia (AMKL; AML M7) accounts for approximately 10% of acute myeloid leukemia (AML) cases in children. In non-Down Syndrome (DS) pediatric patients, this malignancy is particularly aggressive and associated with poor outcomes^1–4^. Oncogenic gene fusions play a primary role in the aberrant activation of oncogenic pathways driving pathogenesis in pediatric patients^5–7^. The CBFA2T3-GLIS2 gene fusion, generated by a chromosome 16 inversion [inv(16)(p13.3q24.3)]^5,8^, is prevalent among pediatric non-DS AMKL patients (between 12-30%) but is not observed in adult AMKL or in other AML subtypes^5,6,8–10^. The prognosis for pediatric AMKL patients harboring CBFA2T3-GLIS2 is significantly worse as compared to patients carrying other gene fusions^5,6,9,11^. However, the molecular mechanisms of CBFA2T3-GLIS2-driven AMKL remain unclear.

CBFA2T3 (also known as ETO2/MTG16/RUNX1T3) has been shown to maintain hematopoietic stem cell (HSC) self-renewal potential and promote AML proliferation^12–14^. In normal myeloid cells, CBFA2T3 acts as a transcriptional corepressor that plays a critical role in the differentiation of erythroblasts and megakaryoblasts, as well as in the development of megakaryocyte-erythrocyte progenitors (MEP)^15–18^. When fused to GLIS2, the N-terminal portion of CBFA2T3 often lacks its myeloid, nervy, and DEAF-1 (MYND) zinc-finger domain, required for transcriptional repression^5,19^. Nonetheless, the CBFA2T3-GLIS2 fusion protein retains the capacity to bind DNA through the five intact zinc-finger domains of GLIS2, allowing CBFA2T3 localization to atypical genomic loci^7^. GLIS2 (also known as NPHP7), a member of the GLI-similar zinc finger family is not normally expressed in hematopoietic cells^20^. GLIS2 regulates Hedgehog (Hh) signaling, and Hh target gene expression is increased in cells expressing CBFA2T3-GLIS2^5,21^. Notably, GLIS2 enhances the self-renewal capacity of HSC, and the C-terminal fusion of GLIS2 to CBFA2T3 allows the former to be expressed at high levels in hematopoietic cells^5^.

No targeted therapies are available for CBFA2T3-GLIS2-driven AMKL^9,22^. Therefore, the development of tractable preclinical models for this leukemia is particularly important^23^. We describe novel murine models of CBFA2T3-GLIS2-driven AMKL that allowed us to identify, for the first time, a critical contribution for the GLIS2 moiety in this disease. Furthermore, we show that a Ras activating mutation promotes CBFA2T3-GLIS2-dependent AMKL by decreasing latency and increasing penetrance. The coordinated modulation of pro- and anti-apoptotic proteins by CBFA2T3-GLIS2 (and by GLIS2 alone), evoking a state of apoptotic priming, further suggests a fundamental sensitivity of AMKL cells to BH3 mimetics. Targeting this vulnerability with navitoclax showed remarkable therapeutic effects on both mouse and human AMKL, suggesting a promising new treatment regimen for AMKL.

## Methods

### Mice

Mice (**Supplemental Table 1**) were purchased from The Jackson Laboratory and maintained in a specific pathogen free animal facility at the Maisonneuve-Rosemont Hospital Research Center; the experiments were approved by the Research Center animal care committee in accordance with the Canadian Council on Animal Care guidelines. Male and female mice were used.

### Transplantation and Leukemia Monitoring

For dual-vector transplantation assays, 1e6 mCherry^+^ fetal liver (FL) cells were sorted on a BD FACSAria III Cell Sorter, mixed or not with WT B6.SJL bone marrow (BM) helper cells, and transplanted intravenously via the tail veins of irradiated (800 cGy) C57BL/6 recipients. For the FL vs BM transplantation cohort, BM cells from 8–12 week old B6.SJL mice were depleted of mature lineage markers using the EasySep™ Cell Separation kit (STEMCELL). Isolated cells were cultured, transduced, and transplanted as described for FL cells. Transduced FL and BM cells were distinguished from host cells by mCherry expression or using antibodies against CD45.1 and CD45.2 (**Supplemental Table 1**). Peripheral blood (PB) was monitored every two weeks by flow cytometry. In addition, mice transplanted with luciferase-expressing vectors were monitored by bioluminescence imaging. 150 mg/kg of XenoLight D-Luciferin (PerkinElmer) was injected intraperitoneally prior to imaging with an IVIS Lumina II LT (PerkinElmer). Luminescence was quantified using LivingImage® software (PerkinElmer), and bioimaging performed until no overt change in luminescence was observed. Mice were sacrificed humanely either when moribund or at the end of the experiment (182 days).

### Retroviral and Lentiviral Constructs

The MSCV-CBFA2T3-GLIS2-IRES-mCherry (CG2) vector was described previously ^5^. The MSCV-IRES-mCherry (MIC) empty vector was generated by replacing CBFA2T3-GLIS2 with an EcoRI/XhoI fragment from MIG. CBFA2T3 (NM_005187.5), GLIS2 (NM_001318918.1), and GLIS2^C265G^ vectors were generated by synthesizing the full-length genes into an empty MIC digested with EcoRI/XhoI (GenScript, Piscataway, NJ, USA). The MSCV-Luciferase-IRES-Nras^G12D^ vector (Nras) was described previously^24^. The MSCV-Luc-IRES (Luc) empty vector was generated by replacing Nras^G12D^ with a 5’-ATC GAT ACC GGT GCG GCC GCA TTA TCG TGT TTT TCA AAG GAA AAC C-3’ BmgBI/ClaI fragment. MSCV-GFP-IRES (MGI) was created by amplifying the GFP sequence from the pMIG vector using primers harboring NcoI/BamHI restriction sites, and then by replacing the luciferase cassette from Luc empty vector by the digested PCR fragment. The Nras^G12D^ cassette from the Nras vector was amplified by PCR using primers harboring NotI/ClaI restriction sites, and the digested PCR fragment was subcloned into the MGI vector. All PCR reactions were carried out using the Q5® High-Fidelity DNA Polymerase (New England Biolabs). pCL-ECO was used to generate retroviral virions. The doxycylin-inducible vectors were created by first blunt subcloning of CBFA2T3-GLIS2 or GLIS2 cDNA into the pENTR/D-TOPO (Invitrogen), followed by Gateway recombination (Thermo Fisher Scientific) into the lentiviral entry vector pCW57.1. sgRNA sequences targeting human AAVS1 (sgAAVS1, 5’-CAC TGT GGG GTG GAG GGG A-3’) and GLIS2 (sgGLIS2_1, 5’-TGG CCG AGG TTT CAA CGC C-3’; sgGLIS2_2, 5’-TGT CAA CGA TTA CCA TGT C-3’) were subcloned in LentiCRISPRv2GFP as previously described^25^. psPAX2 and pMD2.G were used to generate lentiviral virions. See **supplemental Table I** for a list of the vectors used.

### Cell Lines and Cell Culture

HEK293T embryonic kidney cells (ATCC) were cultured in Dulbecco modified Eagle medium (DMEM) supplemented with 10% fetal bovine serum (FBS, GIBCO), 1 mM sodium pyruvate and antibiotics. Human AMKL cell lines M07e, WSU-AML and RS-1 cells were described previously^5^ and were cultured in Roswell Park Memorial Institute medium (RPMI) 1640 media (Wisent Bio Products, St-Bruno, QC, Canada) supplemented with 10% FBS (GIBCO), 10 ng/mL human recombinant IL-3 (BioLegend, San Diego, CA, USA) and antibiotics. All cells were maintained at 37 °C in a humidified incubator with 5% CO2. For short-term *in vitro* culture, PDX cells were maintained in StemSpan™ SFEM II serum-free media supplemented with 10ng/mL recombinant human IL-3, IL-6, thrombopoietin, stem cell factor (SCF) and FLT3L. ABT-199 and ABT-263 were purchased from Selleck Chemicals (Houston, TX, USA) or MedChemExpress (Monmouth Junction, NJ, USA). Bone marrow (BM) cells from leukemias were maintained in RPMI supplemented with 10% FBS, 6 ng/mL recombinant mouse (rm) Il-3, 10ng/mL rmIl-6, and 50 ng/mL rmSCF, and antibiotics. Cytokines were either purchased from STEMCELL Technologies (Vancouver, BC, Canada) or BioLegend (San Diego, CA, USA).

### Fetal Liver Cell Isolation

Timed pregnancies of C57BL/6 and B6.SJL mice were initiated to obtain CD45.1^+^CD45.2^+^ embryos. Fetal livers isolated at embryonic day 13.5-14.5 were gently crushed with a syringe rubber tip over 70 μm nylon mesh and washed twice with PBS supplemented with 2% FBS (Thermo Fisher Scientific). FL cells were seeded at 1e6 cells/mL in DMEM supplemented with 10% FBS, 1 mM sodium pyruvate, 1e-5M β-mercaptoethanol, 6 ng/mL rmIl-3, 10ng/mL rmIl-6, and 50 ng/mL rmSCF for 24h prior to transduction.

### Transfections and Transductions

Transfections of plasmid DNA in HEK293T cells were performed using Lipofectamine® 2000 (Invitrogen) according to the manufacturer’s instructions. Retroviral supernatants were prepared as described ^26^. pCL-ECO and MSCV-based vectors were co-transfected at a 1:1 ratio. FL cells were seeded at 1e6 cells/mL with fresh media (plus cytokines) mixed to an equal volume of viral supernatants, with 6 μg/mL hexadimethrine bromide and spun at 900g for 90 min at room temperature. FL cells were incubated for 48 h prior to experiments. Lentiviral particles were produced by co-transfection of psPAX2, pMD2.G, and transfer vectors in HEK293T cells. Supernatants were collected 48h-60h after transfection.

### RNA-Sequencing and Bioinformatic Analyses

Cells of interest were sorted with a BD FACS ARIA III directly into TRIZOL reagent. Total RNA was extracted to generate transcriptome libraries. 75 cycles of single-end sequencing were performed using an Illumina NExtSeq500. Sequences were trimmed for sequencing adapters and low quality 3’ bases using Trimmomatic version 0.35 and aligned to the reference mouse genome version GRCm38 (or mm10, gene annotation from Gencode version M13, based on Ensembl 88) using STAR version 2.5.1b^27^. Normal murine megakaryoblast data was obtained from the previously published dataset by Dang et al (GEO accession: GSE95081)^28^. Differential gene analysis was performed using R version 3.6.3 (R Project for Statistical Computing) with DESeq2 package version 1.0.12^29^. Heat maps of differentially expressed genes were generated with pheatmap package version 1.0.12. Gene set enrichment analysis (GSEA) was performed using GSEA software version 4.0.3 (Massachusetts Institute of Technology); signal-to-noise ranking criteria was used as a default. ssGSEA^30,31^ was used to derive a signature score for each progenitor myeloid subset as implemented in DRPPM-EASY^32^. Gene set was derived from mRNA profiling of HSC, GMP, CMP, MEP, and MPP as described^28^. RNA sequencing data was mapped by STRONGARM^33^. HTSEQ was used to calculate raw read counts^34^ and normalized to FPKM. LIMMA was applied to derive differentially expressed genes based on an adjusted p-value cutoff of 0.05. See **supplemental Table 3** for the complete gene list.

### Cell Death Assays

Cells were seeded in 96-well, non-adherent round bottom plates at densities of 2.5e5 cells/mL and treated BH3 mimetics for 72h. Drugs were added and used at a final DMSO concentration of 0.5% for human cell lines and primary murine leukemia, and of 0.1% for PDX-derived cells. Cell death was measured using the annexin V-APC Apoptosis Detection Kit (BioLegend) according to the manufacturer’s instructions and analysed with a BD Fortessa X20 (BD) flow cytometer. Percentages of annexin V^+^ cells were normalized, and EC50 were calculated by curve fitting using GraphPad Prism v9.0 software.

### Western Blot

Cells were lysed in 50 mM Tris-HCl (pH 8.0), 150 mM NaCl, 0,1% sodium deoxycholate and 1% NP-40 with Complete EDTA-free protease inhibitors (Roche) and sonicated 10 s on ice. Cell extracts were separated by SDS-PAGE and transferred to a PVDF membrane (Bio-Rad) for immunoblotting. Primary antibodies used are described in **supplemental Table 1**. IgG antibodies conjugated to HRP were detected using the enhanced chemiluminescence (ECL) detection kit (DuPont) and acquired with an Azure c600 imaging system (Azure Biosystems).

### Real-Time Quantitative PCR

Total RNA was isolated in Trizol® reagent according to the manufacturer’s instructions (Invitrogen). The RevertAid H Minus First Strand cDNA Synthesis Kit was used with random hexamer primers (Thermo Fisher Scientific) to reverse transcribe 1 μg of total RNA in a final volume of 20 μl. Real-time quantitative PCR (RT-qPCR) reactions were performed in 96-well plates using 5ng of cDNA samples using the Luna Universal qPCR Master Mix (New England Biolabs) according to the manufacturer’s instructions. Amplification levels were detected using the QuantStudio 12K Flex Real-Time PCR System (Thermo Fisher Scientific) programmed to run at 95 °C for 1 min before starting 40 cycles of 15 s at 95 °C and 30 s at 60 °C. The reactions were run in triplicate and the quantification performed using average Cts values. The ΔΔCT method was used to determine relative quantities of target genes. Primers used for RT-qPCR reaction are described in **supplemental Table 1.**

### Histology

Hematoxylin-eosin stains were performed according to standard procedures. Immunohistochemistry reactions were performed on the Bond RX Stainer (Leica Biosystems, Buffalo Grove, IL, USA) according to the manufacturer’s instructions using an anti-Factor VIII antibody (Biocare Medical, Concord, CA, USA). Detection of tissue-bound primary antibody was performed using the Bond Intense R Detection System (Leica Biosystems). The stained slides were converted to digital data using the NanoZoomer Digital Pathology 2.0-HT digital slide scanner (Hamamatsu).

### Patient-Derived Xenografts

Primary AML blasts from two pediatric patients (M7012 and M7014) were obtained with patient or parent/guardian-provided informed consent under protocols approved by the Institutional Review Board at St. Jude Children’s Research Hospital. Cryo-conserved patient cells were thawed and i.v. injected into 8–week old NSG-SGM3 recipient mice for expansion. Whole BM and spleen cells were processed to obtain a single-cell suspension and erythrocytes were lysed with ACK buffer (150 mM NH4Cl, 10mM KHCO3, 100 μM Na2EDTA, pH 7.4) for 5 minutes. Cells were incubated with anti-mouse CD45 MicroBeads and subjected to negative depletion (Miltenyi Biotec Inc). Cells were either used fresh for *ex vivo* experiments or preserved in CryoStor® CS10 freezing media (STEMCELL Technologies). For *in vivo* experiments, 1-2e6 cells derived from patient M7012 were i.v. injected into 8–week old NSG-SGM3 female recipient mice, as previously described. Peripheral blood analysis was performed 2 weeks posttransplantation to monitor disease progression, represented by the presence of >2% mCD45^-^ hCD56^+^ blasts as assessed by flow cytometry. 50mg/kg ABT-263 was dissolved in gavage media consisting of 60% phosal 50 PG (Thermo Fisher Scientific), 30% poly(ethylene glycol) 400 (MilliporeSigma), and 10% ethanol, and administered *per os* 5 times per week starting 18 days post-transplantation, over the course of 3 weeks. Gavage media was used as vehicle control. Mice were sacrificed humanely when moribund. Human blast infiltration was evaluated by the presence of mCD45^-^hCD56^+^ blasts, either upon sacrifice or 39 days post-transplantation, whichever occurred first.

### Proteomic Profiling of Pediatric Patients Samples

100 μg of protein from normal primary megakaryoblasts and CBFA2T3-GLIS2 patient samples was analyzed in technical replicate by tandem mass tag (TMT) mass spectrometry at the Center for Proteomics and Metabolomics at St. Jude Children’s Research Hospital. Protein lysates were digested and desalted and then underwent 16-plex TMT labeling. Samples were analyzed for total proteome by low pH reverse-phase liquid chromatography tandem mass spectrometry. Retrieved data was searched against the UniProt human database to identify peptides. This was determined by assigning related mass addition to all possible amino acid residues followed by an algorithm that evaluates the accuracy of modification localization. Peptide quantification data was corrected for mixing errors and summarized to derive protein quantification results.

### Flow Cytometry Analysis

Cells were stained with Zombie-Aqua Fixable Viability dye (BioLegend) following manufacturer’s instructions followed by extracellular staining in PBS 2%FBS. Antibodies used are described in **Supplemental Table 1**. Samples were run on a BD Fortessa X20 flow cytometer and data analyzed with FlowJo software (BD).

### Statistical Analyses

Statistical significance relative to control conditions was established by one-way ANOVA followed by Tukey’s multiple comparison test for multiple groups unless specified otherwise.

### Data availability

Material and reagents generated during the current study are available from the corresponding authors on reasonable request. RNA sequencing data is uploaded to the Gene Expression Omnibus (GEO) data repository (GSE208187).

## Results

### Mouse Models of CBFA2T3-GLIS2-Driven AMKL Recapitulate Human Disease

Pediatric AMKL patient samples harbor relatively few genetic mutations but display high gene copy number alterations^5,35^. Despite the generally low frequency of hotspot mutations cooccurring with CBFA2T3-GLIS2, the presence of oncogenic *NRAS* mutations has been reported^6^. Furthermore, mutations in the JAK-STAT pathway, which can activate RAS-MAPK signaling^36^, are found in megakaryoblastic cell lines, DS- and non-DS AMKL patients, and CBFA2T3-GLIS2-positive patients^5,6,37,38^. Given that NRAS is frequently activated in human AML^39^, and since previous mouse models of AML fusions co-expressing mutant Nras exhibited enhanced penetrance and aggressiveness of the disease^24^, we reasoned that oncogenic NRAS would promote CBFA2T3-GLIS2-driven leukemogenesis in mice.

To test the capacity of mutant NRAS to enhance the leukemogenicity of CBFA2T3-GLIS2, we used a “mosaic” strategy allowing concomitant expression of oncogenes and reporters^24,40^. Different combinations of bicistronic retroviral vectors encoding (i) mCherry alone (MIC) or with CBFA2T3-GLIS2 (CG2)^5^, and (ii) luciferase (Luc) alone or with constitutively active Nras^G12D^ (Nras)^24^, were used to transduce E13.5-14.5 FL-derived hematopoietic stem and progenitor cells (HSPC) (**Fig. 1A**). Lethally irradiated wild-type (WT) mice were then reconstituted with sorted mCherry^+^ FL cells and monitored for disease onset by PB immunophenotyping and bioluminescent imaging for up to 182 days (**Fig. 1B**). FL cells expressing CG2 and empty luciferase vector (CG2-Luc) efficiently induced leukemia in 40% of transplanted mice within this time period (**Fig. 1C**), consistent with recent reports proposing that expression of the CBFA2T3-GLIS2 fusion is sufficient to induce leukemia^7,9^. When compared with empty luciferase vector (CG2-Luc), the co-expression of oncogenic Nras (CG2-Nras) significantly increased the penetrance of leukemia (65% vs 40%), with mice showing a median survival of 74 days (vs >182 days for CG2-Luc) (**Fig. 1C**). Notably, CG2-Nras mice manifested a marked increase in luciferase signal over time, suggesting positive selection of leukemic blasts expressing oncogenic Nras (**Fig. 1D, Supplemental Fig. 1A**). Mice receiving CG2-Luc and CG2-Nras-expressing FL cells displayed pronounced splenomegaly with variable percentages of mCherry^+^ cells in the BM (**Fig. 1E**). The immunophenotype of mCherry^+^ BM cells from moribund CG2 recipient mice was indicative of AMKL; mCherry^+^ cells from mice that received CG2-transduced FL cells harbored a higher percentage of BM cells co-expressing CD41 and CD61 and immature megakaryocyte progenitors (c-Kit^+^CD41 ^+^) when compared to engrafted mCherry^+^ cells from control mice (MIC-Luc and MIC-Nras). Moreover, BM cells from CG2-Luc and CG2-Nras mice contained a lower percentage of cells expressing lineage markers Mac-1/Gr-1 (**Fig. 1F-G**), B220, CD3, and TER119 (**Supplemental Fig. 1B**), indicating a skewing towards megakaryocytic differentiation. In addition, leukemic blasts isolated from the spleen of CG2-Luc and CG2-Nras mice displayed a significant increase in CD41 without enrichment of macrophage (Mac-1) or granulocyte (Gr-1) lineages (**Supplemental Fig. 1C**). Interestingly, most of the mCherry^+^ BM cells from CG2 recipients lacked CD45 expression (**Supplemental Fig. 1D**), characteristic of the “RAM phenotype” observed in human leukemias with the CBFA2T3-GLIS2 fusion^9,41^. Although some control mice (MIC-Luc, MIC-Nras) developed different forms of leukemia, most showed enrichment of Mac-1 and Gr-1 expressing BM cells, and no control mice developed AMKL (**Fig. 1C, F-G**). To further define the clinical characteristics of CG2-derived murine leukemias, we performed histological analysis of formalin-fixed and paraffin embedded spleen sections from CG2 recipients revealing highly altered follicular architecture, infiltration of monomorphic cells, and increased Factor VIII staining (**Fig. 1H**). Abnormally high numbers of white blood cells were observed in PB smears from CG2 recipients (**Fig. 1H**).

**Figure 1.**
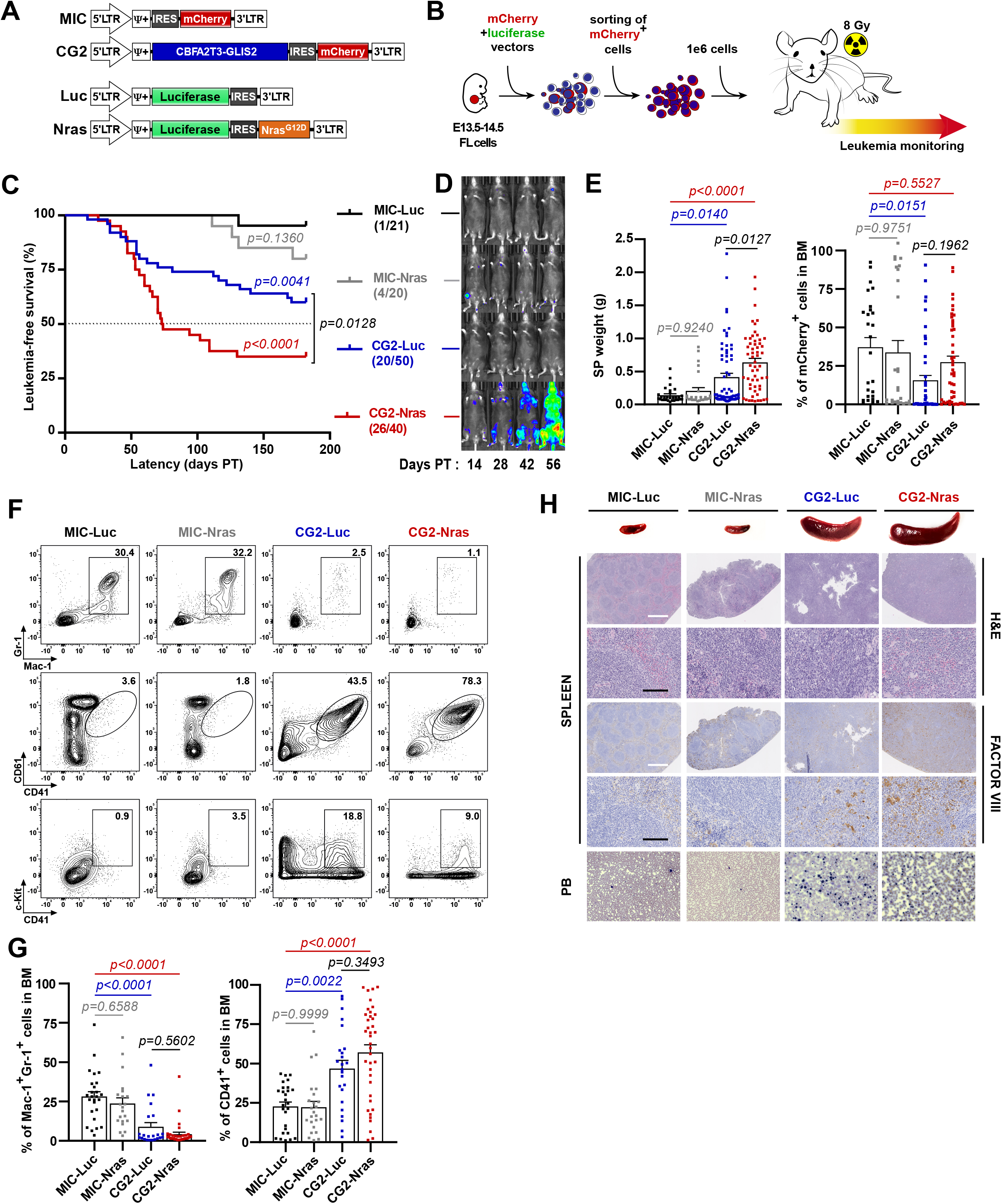
Mosaic AMKL Mouse Models Phenocopy Pediatric Human AMKL. **A.** MSCV-based retroviral vectors were designed to be co-transduced in mouse FL cells. The mCherry fluorescent protein construct expresses CBFA2T3-GLIS2 or not, and the *Photinus pyralis* “firefly” luciferase construct co-expresses Nras^G12D^ or not. **B.** CD45.1/45.2-double positive FL cells were co-transduced with a combination of mCherry and luciferase constructs, and mCherry-expressing cells were sorted 48h post-transduction. Lethally irradiated (8Gy) mice were transplanted with 1e6 mCherry^+^ cells and subsequently monitored for AMKL markers. **C.** Kaplan-Meier plot showing the percentage of leukemia-free survival from each vector combination group over time. Numbers represent leukemic mice over the total number of mice in each group. Statistical significance was measured by the log-rank test (Mantel-Cox). **D.** Bioluminescent imaging of primary recipient mice 14-, 28-, 42- and 56-days following transplantation; representative images. **E.** Spleen weight and percentage of mCherry^+^ cells recovered from the BM at time of sacrifice. **F, G.** Representative flow cytometry plots (**F**) and quantification of flow cytometry markers (**G**) from mCherry^+^ BM cells at the time of sacrifice. **H.** Representative images of spleen from MIC controls and CG2 leukemic mice with corresponding spleen sections stained with hematoxylin and eosin or anti-factor VIII antibody. Black scale bar = 1mm (magnification 2.5x), white scale bar = 100μm (magnification 20x). PB smears (original magnification 1000x) stained with Wright-Giemsa.

The rationale for using FL cells to generate our models was based on the idea that due to the early disease onset observed in pediatric CBFA2T3-GLIS2 AMKL patients, the oncogenic fusion might require hematopoietic cells of fetal origin to drive transformation^42^. To assess whether BM HSPC could reproduce the leukemias obtained with FL cells, we transplanted lineage depleted BM or FL cells transduced with CG2. CG2 expression in either BM HSC or FL HSPC led to a moderately penetrant leukemia with CD41^+^ and CD61^+^ BM blasts (**Supplemental Fig. 1E**-**F**), suggesting that BM cells can also be used to generate murine models of CBFA2T3-GLIS2 AMKL. Altogether, our data show that, as opposed to a previously published transgenic CG2-expressing model where most of the leukemia harbored an AML-like phenotype (Mac-1^+^Gr-1^+^)^10^, retrovirus-mediated CG2 expression in mouse HSPC consistently induce a penetrant leukemia phenotypically similar to human AMKL (CD41 ^+^Mac-1 ^-^Gr-1 ^-^). In addition, the coexpression of CG2 and oncogenic Nras increases leukemia aggressiveness by raising penetrance and decreasing latency without overtly affecting the AMKL phenotype, emphasizing its usefulness as a model of human AMKL, and enabling the identification of cooperating oncogenes.

### CG2-Driven Murine Leukemias Correlate with the Human AMKL Transcriptome

We next evaluated the pertinence of using these models to elucidate molecular pathways involved in the human pathology by performing transcriptomic analyses of FL cells transduced with either CG2 or empty MIC vector. Within 2 days of transduction with CG2, the gene expression pattern of FL cells already displayed significant similarities to that of CBFA2T3-GLIS2-expressing AMKL from different cohorts of pediatric patients (**Fig. 2A**, **Supplemental Fig. 2A** and **Supplemental Table 2**)^5,6,9^. Since the overall clinical and phenotypic characteristics of CG2-driven leukemias were equivalent in the presence or not of constitutively-activate Nras (**Fig. 1**), we compared gene expression signatures from CG2-Luc and CG2-Nras leukemic cells to determine whether more subtle molecular differences could be observed. Unsupervised clustering revealed analogous transcriptional signatures between the CG2-Luc and CG2-Nras leukemias in comparison to control MIC-Luc BM cells (**Fig. 2B**, **Supplemental Fig. 2B** and **Supplemental Table 2**). We then probed whether the presence of oncogenic Nras was affecting gene signatures related to human AMKL. We observed that both CG2-Luc and CG2-Nras leukemias showed a gene signature consistent with markers of human CBFA2T3-GLIS2-driven leukemia, as well as with megakaryoblast and stemness markers (**Fig. 2C**). Furthermore, gene expression profiles of CG2-driven leukemias showed enrichment of a previously published expression signature from a transgenic mouse model of CBFA2T3-GLIS2 leukemia (**Fig. 2D**, **Supplemental Fig. 2C**)^10^, and correlated also with the gene profiles of AMKL patients as compared to AML (**Fig. 2E**)^43^. Single-sample gene set enrichment analysis (ssGSEA) confirmed that the gene signatures of CG2^+^ AMKL cells were consistent with MEP and common myeloid progenitor (CMP) signatures (**Fig. 2F** and **G)**, while MIC-transduced BM cells were enriched with other lineages of multipotent progenitors (MPP), granulocyte/macrophage progenitors (GMP), and HSC signatures (**Supplemental Fig. 2D, Supplemental Table 3**)^28^, thus confirming the AMKL-like transcriptome inherent to the CG2-derived leukemias. Altogether, these results indicate that retroviral-based mosaic mouse models of CG2^+^ AMKL faithfully replicate human AMKL both phenotypically and transcriptionally, thereby providing a novel, robust platform to dissect the molecular mechanisms and potential therapeutic vulnerabilities of this disease.

**Figure 2.**
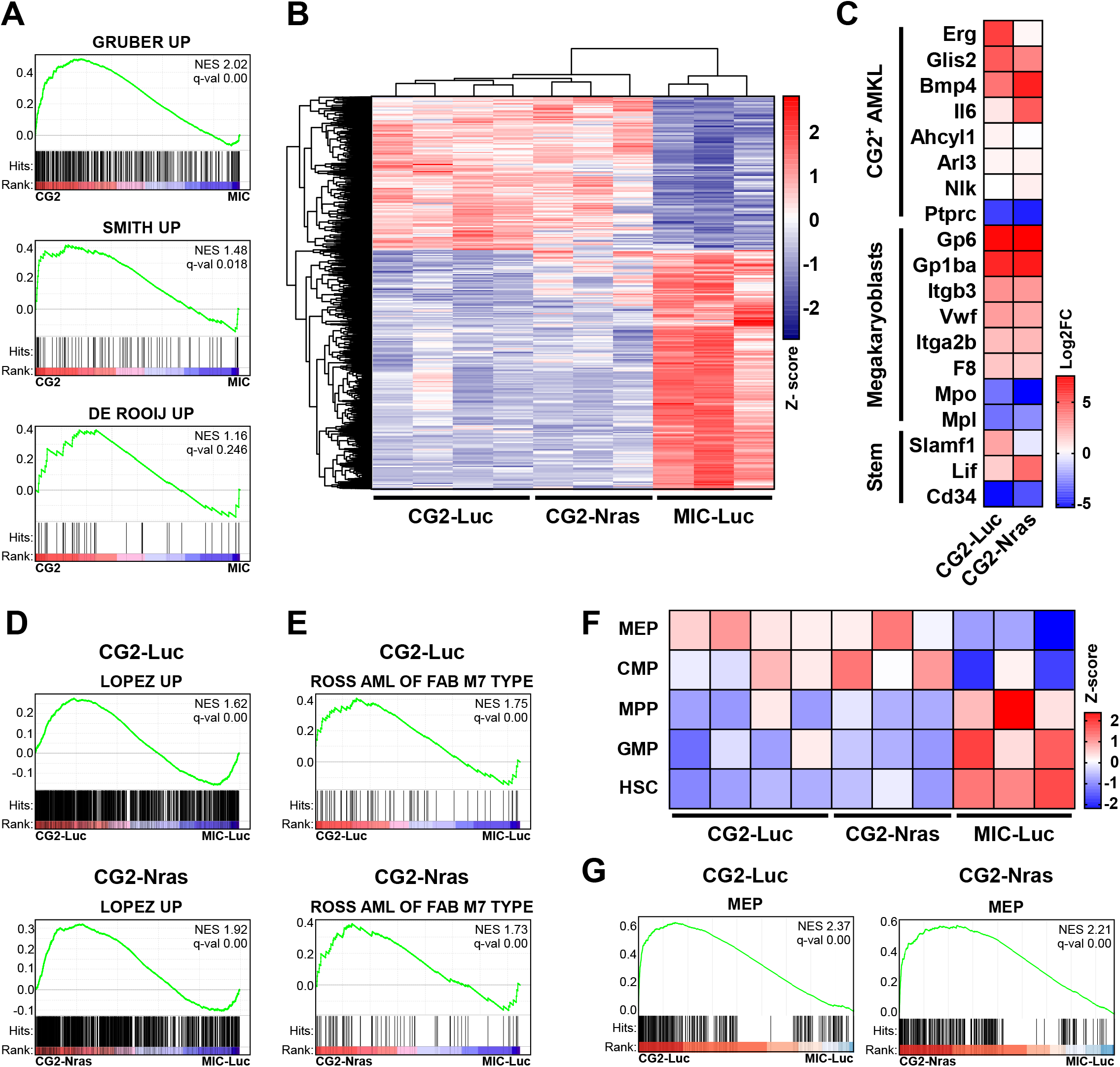
Transcriptional Profiles of CBFA2T3-GLIS2-Induced Murine Leukemias Correlate with Human AMKL. **A.** GSEA of differential gene expression in FL cells expressing CBFA2T3-GLIS2 (CG2) vs control (MIC) for gene sets comprising genes upregulated in CBFA2T3-GLIS2^+^ patients vs fusion-negative AMKL^5,6^ or AML^9^. FL cells were sorted for mCherry^+^cKit^+^Lin^-^ (Gr-1 ^-^, B220^-^, CD3^-^, Ter119^-^, Mac-1+) expression. **B.** Unsupervised hierarchical clustered heatmap of differentially expressed genes (5000 most significant *p*-values; Z-score) in mCherry^+^-sorted BM cells from MIC-Luc, CG2-Luc and CG2-Nras mice at sacrifice. MIC-Luc was used as a control. **C.** Heatmap of differential expression data from (**B**) of genes that are differentially expressed in patients with CBFA2T3-GLIS2 AMKL as well as markers indicative of megakaryoblasts and hematopoietic stemness. **D, E.** GSEA of differential expression data from (**B**) for gene sets containing genes upregulated upon doxycycline induced expression of CBFA2T3-GLIS2 in FL HSPC vs WT BM HSC (**D**)^10^ and signature genes from AMKL (AML-M7; FAB classification) vs other AML subtypes (**E**)^43^. **F.** Heatmap of myeloid lineage signature scores of data from (**B**), calculated by single-sample GSEA from previously published gene signatures from flow-sorted mouse hematopoietic stem cell (HSC), granulocyte-monocyte progenitors (GMP), multipotent common myeloid progenitor (CMP), megakaryocyte-erythroid progenitor cell (MEP), and multipotent progenitors (MPP)^28^. **G.** Corresponding GSEA from (**F**) showing enrichment in MEP signature from CBFA2T3-GLIS2^+^ leukemias.

### CBFA2T3-GLIS2-Positive Cells Exhibit Features of RAS-MAPK Pathway Activation

We showed that activated RAS, which is known to cooperate with other leukemia-associated fusion proteins to promote AML^24,44^, increases penetrance and decreases latency of CG2-dependent AMKL (**Fig. 1**). In light of our findings that CG2 expression is sufficient to generate AMKL, we further examined whether this oncogenic fusion could modulate a transcriptional signature associated with RAS-MAPK pathway activation. To enable the sorting of Nras^G12D^-expressing FL cells, we replaced the luciferase cassette with a GFP cassette to generate a GFP-IRES-Nras^G12D^ bicistronic vector (along with a corresponding GFP-IRES-empty vector; MGI) (**Supplemental Fig. 3A**). We retrovirally co-transduced FL cells with mCherry and GFP vectors, encoding or not CG2 and Nras^G12D^, respectively, and then performed RNA-seq on mCherry^+^GFP^+^Kit^+^lin^-^ (lin^-^ defined as Gr1^-^CD3^-^B220^-^Ter119^-^) sorted cells. Consistent with analyses of the CG2^+^ murine leukemias expressing or not oncogenic Nras, CG2-transduced FL cells presented a similar gene expression signature in the presence or absence of Nras^G12D^ (**Fig. 2B, Supplemental Fig. 3B** and **C**). Furthermore, in both cases, GSEA revealed a transcriptional signature consistent with RAS-MAPK pathway activation (**Supplemental Fig. 3D**)^45^, suggesting that CBFA2T3-GLIS2 aberrantly contributes to the stimulation of this oncogenic pathway.

### GLIS2 Cooperates with Oncogenic Ras to Promote AMKL

To determine the relative contribution of each partner of the CBFA2T3-GLIS2 fusion to the CG2-associated transcriptional signature, FL cells transduced with full-length human CBFA2T3, GLIS2, a DNA-binding deficient mutant of GLIS2 (GLIS2^C265G^)^46^, or CG2 were subjected to transcriptomic analyses. Unsupervised clustering of differentially expressed genes revealed striking similarities between the CG2 and GLIS2 transcriptomes (**Fig. 3A** and **B**). Only a few genes were consistently modulated by both CG2 and CBFA2T3 in FL cells, while GLIS2^C265G^ had a limited effect on gene expression and clustered with empty MIC vector (**Fig. 3A** and **B**). Transcriptional signatures of CG2- and GLIS2-expressing cells both revealed a significant over-representation of pathways involved in oncogenesis (*e.g*., IL2-STAT5 signaling, IL6-JAK-STAT3 signaling, apoptosis) (**Fig. 3C**)^45,47^. Due to strong similarities between the transcriptional programs triggered by CG2 and GLIS2, we tested the potential of GLIS2 to promote AMKL *in vivo*. The ability of GLIS2 to promote AMKL *in vivo* has not yet been evaluated. Of note, previous studies showed that GLIS2 enforces a megakaryocytic phenotype in murine HSPC but promoted only moderate *in vitro* self-renewal capacity as compared to CBFA2T3-GLIS2^5,7^. Thus, we assumed that GLIS2 alone would be insufficient to induce AMKL. Since oncogenic Ras increased the penetrance of CG2-driven AMKL without altering the phenotype, we hypothesized that Nras^G12D^ may cooperate with GLIS2 to induce AMKL *in vivo*. FL cells expressing CBFA2T3, GLIS2, or both GLIS2 and Nras^G12D^ were transplanted into lethally irradiated WT recipients and monitored for leukemia onset. CBFA2T3 alone provoked an aggressive and penetrant leukemia reminiscent of an immature AML phenotype; mCherry^+^ BM cells predominantly expressed Mac-1 and Gr-1 but not CD41 (100% penetrance; **Fig. 3D** and **E**). While GLIS2 alone could not induce leukemia, concomitant expression of GLIS2 and Nras^G12D^ induced CD41^+^ AMKL (33% penetrance) (**Fig. 3D** and **E**), and leukemic GLIS2-Nras mice had enlarged spleens (**Fig. 3F**). These data suggest that the GLIS2-dependent transcriptional program contributes to the AMKL disease phenotype. Notably, GLIS2 recipients that did not develop leukemia also displayed less than 1% chimerism in the BM 6 months post-transplantation (**Fig. 3F**), suggesting either a poor engraftment of GLIS2-expressing FL cells, or lack of sustained proliferation of engrafted cells. Interestingly, PB cells expressing GLIS2 (with or without Nras^G12D^) displayed increased megakaryocytic lineage markers CD41 and CD61, similar to those expressing CG2 25 days post-transplantation (**Fig. 3G**). Taken together, these data suggest a significant contribution from GLIS2 in the transcriptome and phenotype of CBFA2T3-GLIS2-driven AMKL.

**Figure 3.**
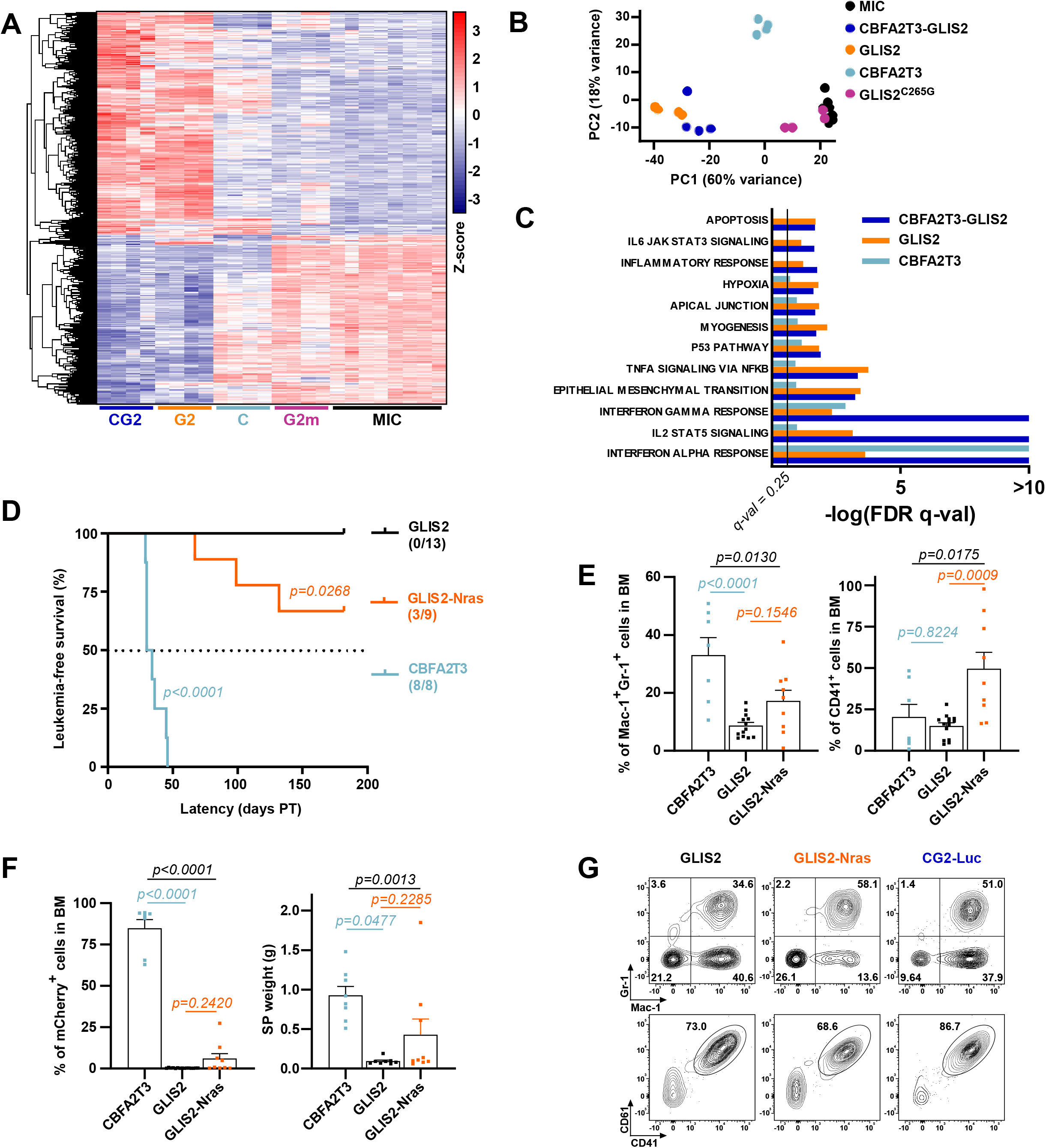
GLIS2 Cooperates with Oncogenic Ras and Drives the Megakaryocytic Identity of Leukemic Blasts. **A.** Unsupervised hierarchical clustered heatmap of differentially expressed genes (5000 most significant *p*-values; Z-score) upon expression of CBFA2T3-GLIS2 (CG2), full-length GLIS2 (G2), full-length CBFA2T3 (C), and GLIS2^C265G^ (G2m) compared to MIC empty vector control in FL cells. FL cells were sorted for mCherry^+^cKit^+^Lin^-^ (Gr-1^-^, B220^-^, CD3^-^, Ter119^-^, Mac-1^+^) expression. **B, C.** Principal component analysis (PCA) of the gene expression profiles (**B**) and bar graph representing hallmark gene sets^45^ significantly overlapping with gene signatures from RNA-seq data described in (**A**). **D.** Kaplan-Meier plot showing the percentage of leukemia-free survival from CBFA2T3, GLIS2 or GLIS2-Nras recipients over time. Numbers represent the leukemic mice over the total number of mice in each group. Statistical significance was measured by the log-rank test (Mantel-Cox) using GLIS2 as a control. **E.** Quantification of flow cytometry markers from mCherry^+^ BM cells at time of sacrifice. **F.** Spleen weight and percentage of mCherry^+^ cells recovered in BM at the time of sacrifice. **G.** Representative flow cytometry histograms plots of markers from mCherry^+^ PB cells 25 days after transplantation.

### Pro- and Anti-Apoptotic Gene Modulation by CBFA2T3-GLIS2 and GLIS2

Among the gene sets controlled by CBFA2T3-GLIS2 and GLIS2 (**Fig. 3C**), genes associated with hallmarks of apoptosis were altered in HSPC expressing either CBFA2T3-GLIS2 or GLIS2 (**Fig. 4A**). Transcript levels of numerous apoptosis-related genes in FL cells were regulated upon expression of either CBFA2T3-GLIS2 or GLIS2, but not following expression of GLIS2^C265G^, CBFA2T3 or empty vector (MIC). More specifically, both GLIS2- and CBFA2T3-GLIS2-expressing FL cells, but not CBFA2T3- or GLIS2^C265G^-transduced cells, displayed increased expression of anti-apoptotic *Bcl2, Bcl2l1* (Bc1-x_L_), *Bcl2l2* (Bcl-w) and *Mcl1*, and of pro-apoptotic *Bim, Bik, Bok*, and *Noxa1* (**Fig. 4B**). Overall, a pronounced overlap in the apoptotic gene signature exists between FL cells expressing CBFA2T3-GLIS2 and GLIS2, but not CBFA2T3, suggesting that such genes might be transcriptional targets of GLIS2 (**Fig. 4B**). In addition to their transcriptional regulation, higher protein levels of Mcl1, Bcl-2, Bcl-x_L_ and Bim was observed in FL cells expressing CBFA2T3-GLIS2 (**Fig. 4C**). To evaluate whether CG2-driven AMKL exhibits a similar overall increase in apoptosis-related gene expression, we isolated BM from sick CG2-Luc and CG2-Nras mice and sorted premature leukemic megakaryoblasts for RNA-seq analysis. CG2-dependent AMKL, in the presence or absence of activated Ras, showed increased expression of pro- and anti-apoptotic genes as compared to WT premature megakaryoblasts (**Fig. 4D**). Anti-apoptotic *Bcl2l1* and *Bcl2* expression was generally increased in CG2-expressing blasts when compared to normal WT blasts. The dysregulation of pro- and anti-apoptotic proteins in primary human AMKL samples was also investigated. We used a proteomic approach to assess BCL-2 family members in CBFA2T2-GLIS2^+^ AMKL, which revealed increased protein levels of anti-apoptotic BCL-2 in CBFA2T3-GLIS2^+^ pediatric AMKL as compared to normal megakaryoblasts (**Fig. 4E**). Furthermore, transcript levels of *BCL2* are significantly upregulated in CBFA2T3-GLIS2^+^ patient samples as compared to AMKL driven by other fusion proteins (**Fig. 4F**), suggesting that BCL-2 is preferentially increased in patients harboring CBFA2T3-GLIS2, but does not constitute a general hallmark of pediatric AMKL.

**Figure 4.**
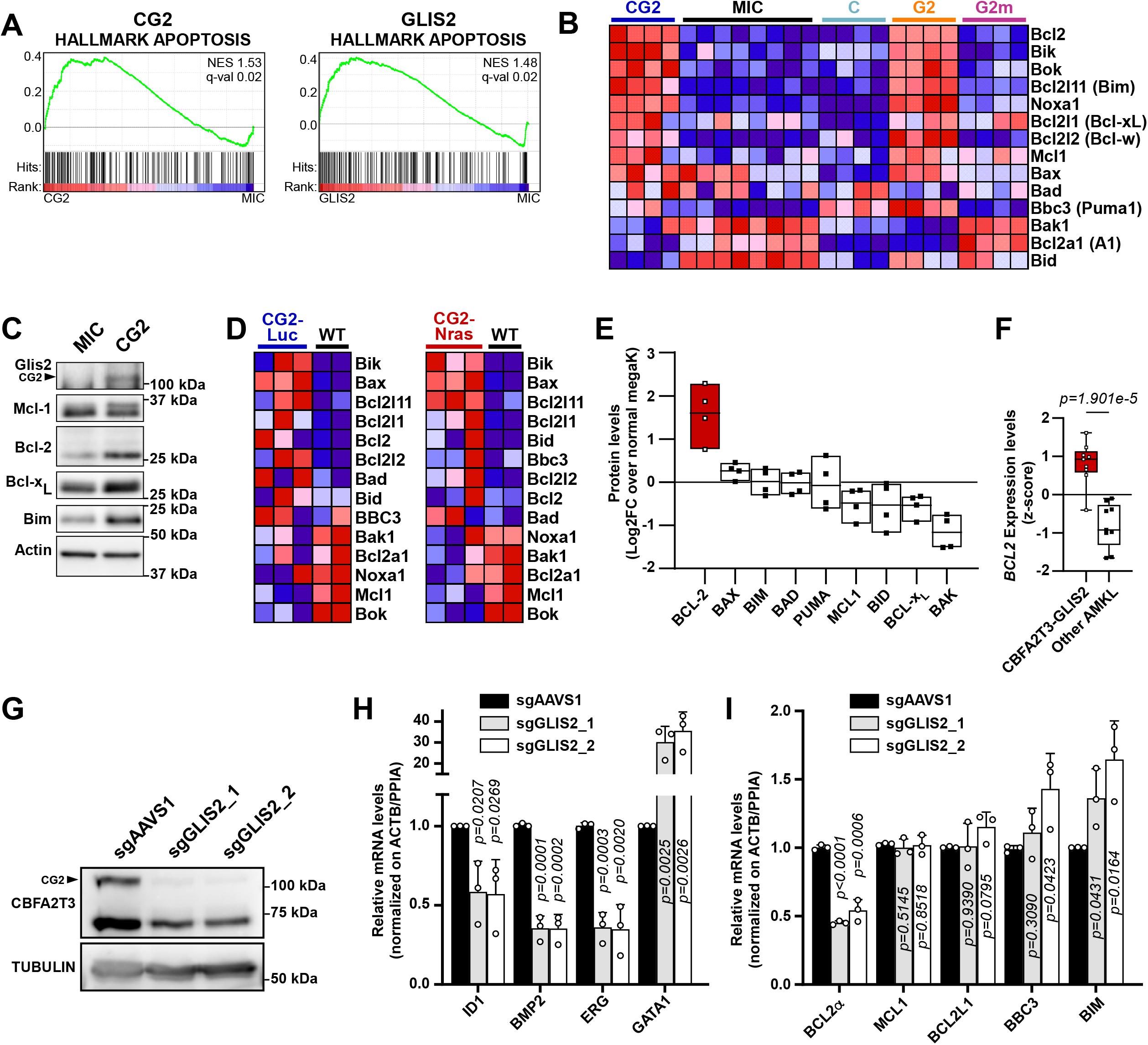
Regulation of Pro- and Anti-Apoptotic Gene by GLIS2 and CBFA2T3-GLIS2. **A.** GSEA plot of expression data obtained from FL cells transduced with CBFA2T3-GLIS2 or GLIS2 showing enrichment of a published apoptosis gene signature^45^. **B.** Heatmap of differentially expressed genes related to apoptosis from RNA-seq data from FL cells expressing CG2, GLIS2, CBFA2T3, and GLIS2^C265G^ compared to MIC empty vector control. Increased expression (red), decreased expression (blue). **C.** FL cells were transduced with empty MIC or CBFA2T3-GLIS2 virions and incubated for 48h. Total cell lysates were immunoblotted with anti-Glis2, anti-Mcl1, anti-Bcl-2, anti-Bcl-xL and anti-Bim antibodies. Anti-pan-actin antibody was used to control for equivalent loading. Representative of two independent experiments. **D.** Heatmap of differentially expressed genes related to apoptosis from BM cells of leukemic CG2-Luc or CG2-Nras mice. Cells were sorted for mCherry^+^CD41^+^cKit^+^Lin^-^ (Gr-1^-^, B220^-^, CD3^-^, Ter119^-^, Mac-1^+^) expression 48h post-transduction; RNA was extracted and subjected to genome-wide RNA sequencing. WT controls consist of C57BL/6 BM cells sorted for CD41^+^cKit^+^Lin^-^ (Gr-1^-^, B220^-^, CD3^-^, Ter119^-^, Mac-1^+^). Increased expression (red), decreased expression (blue). **E.** Relative levels of apoptosis-related proteins between patient samples harboring the CBFA2T3-GLIS2 fusion and normal megakaryoblasts (megaK). Each sample had two replicates and proteins were quantified by tandem mass tag labeling and mass spectrometry. **F.** Expression levels of *BCL2* from pediatric AMKL patients with CBFA2T3-GLIS2 or other genetic subtypes of AMKL. Data from GSE35203 ^5^. **G.** M07e cells were transduced with virions encoding GFP, hCas9, and sgRNAs targeting the AAVS1 locus or GLIS2 (sgGLIS2_1 and _2). Cells were sorted for GFP expression 7-8 days after transduction, and total cell lysates were immunoblotted with anti-CBFA2T3 antibody. Anti-α-tubulin was used to control for equivalent loading. Representative of three independent experiments. **H.** Expression levels of putative transcriptional targets of CBFA2T3-GLIS2 and of (**I**) apoptosis genes were quantified by realtime qPCR in M07e cells from (**G**). Values were normalized to the geometric mean of *ACTB* and *PPIA* values. Data are shown as mean ± s.d. (n = 3).

GLI1 and GLI2 are reported to induce the transcription of *BCL2* through promoter activation^48–50^, and CBFA2T3-GLIS2 may bind to the super-enhancer region of *BCL2*^7^. As such, we hypothesized that CBFA2T3-GLIS2 could regulate *BCL2* in AMKL cells. To investigate whether endogenous CBFA2T3-GLIS2 modulates the expression of *BCL2*, we designed specific guide RNAs (sgRNA) targeting GLIS2 to achieve CRISPR-mediated gene knock-out in CBFA2T3-GLIS2-expressing AMKL cell lines. Two different sgRNAs significantly decreased the levels of endogenous CBFA2T3-GLIS2 protein in M07e AMKL cells (**Fig. 4G**). In addition, CRISPR-mediated depletion of CBFA2T3-GLIS2 resulted in reduced mRNA levels of (i) known GLIS2 transcriptional target *BMP2*^51^, and also of (ii) putative CBFA2T3-GLIS2 targets *ID1* and *ERG* (**Fig. 4H**)^5,7^. Strikingly, *GATA1* expression, a gene involved in megakaryocyte homeostasis and negatively regulated by CBFA2T3-GLIS2^7^, was increased >20-fold following GLIS2-specific sgRNA expression, further confirming the impairment of CBFA2T3-GLIS2 function (**Fig. 4H**). Moreover, reduced expression of *BCL2* (*BCL2α*), with a concomitant increase in *BBC3* (*PUMA*) and *BCL2L11* (*BIM*) levels was observed upon CRISPR-mediated depletion of endogenous CBFA2T3-GLIS2 (**Fig. 4I**). Altogether, these results suggest that CBFA2T3-GLIS2 and/or GLIS2 triggers a transcriptional program that might prime AMKL blasts for apoptosis.

### CBFA2T3-GLIS2-Expressing Cells Are Sensitive to BH3 Mimetics

Human and murine cells expressing CBFA2T3-GLIS2 express higher levels of anti-apoptotic BCL-2 as well as pro-apoptotic BAX, BIM and BAD (**Fig. 4A-F**). This profile is consistent with a cellular state of apoptotic priming, during which cells might be sensitive to BH3 mimetics^52,53^. To assess this possibility, we treated human CBFA2T3-GLIS2^+^ cell lines with venetoclax (ABT-199) or navitoclax (ABT-263), specific inhibitors of BCL-2 or both BCL-2 and BCL-x_L_, respectively^54,55^. M07e and RS-1 cells were unaffected by ABT-199(EC_50_>10μM), but sensitive to ABT-263 (EC_50_= 0.80 and 0.94μM, respectively), suggesting that BCL-x_L_ and BCL-2 are both key players modulating survival in these cells (**Fig. 5A, B**). WSU-AML cells showed striking sensitivity to both inhibitors (EC_50_<150nM)(**Fig. 5A, B**).

**Figure 5.**
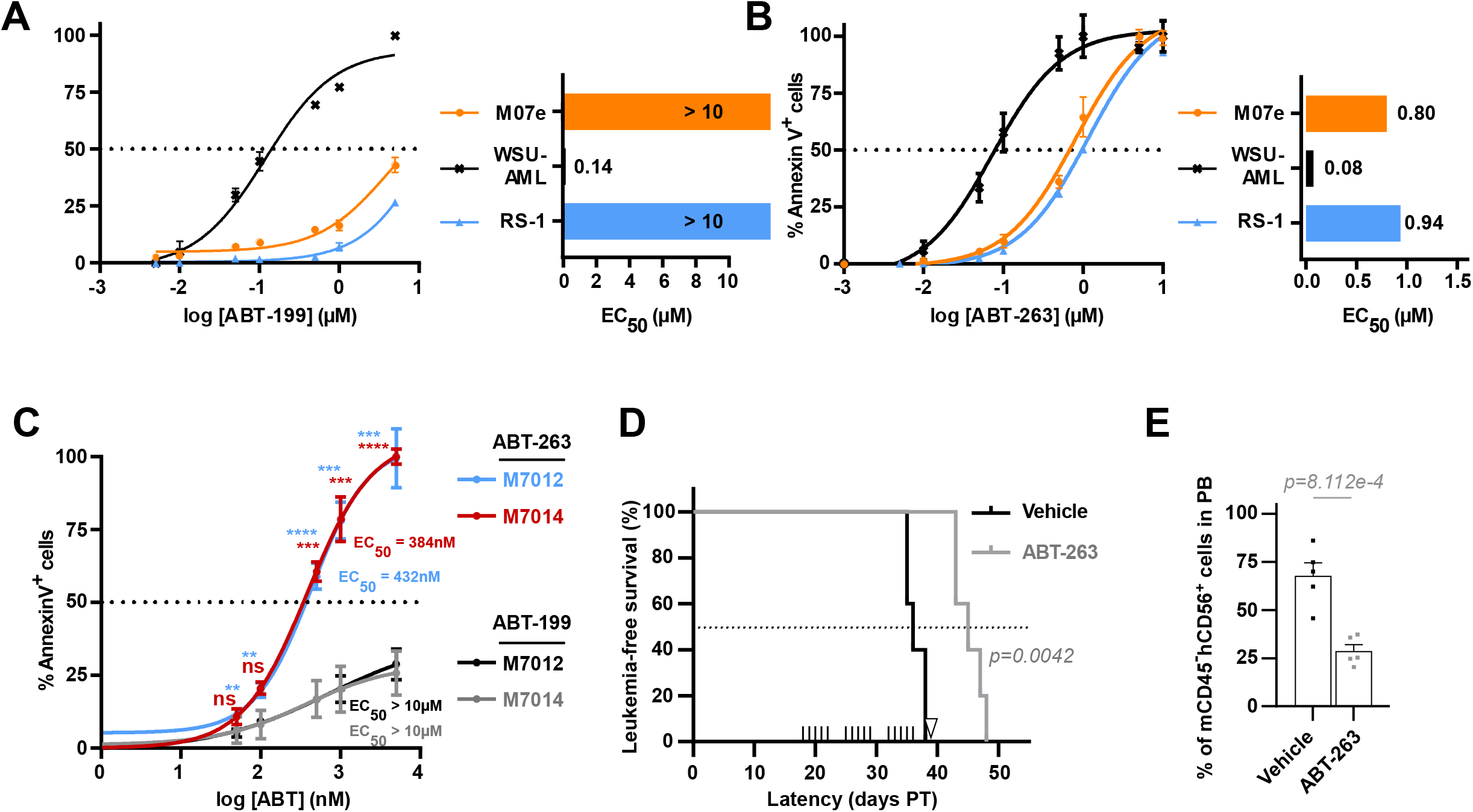
CBFA2T3-GLIS2-Expressing Leukemic Cells Are Sensitive to BH3 Mimetics. Cellular apoptosis and EC50 values for AMKL cell lines (M07e, RS1 and WSU-AML) treated with ABT-199 (**A**) or ABT-263 (**B**) at increasing doses for 72h. Data are shown as mean ± s.d. (n = 3). **C.** Dose-response of PDX-derived cells from two different pediatric patients (M7012 and M7014) treated 72h with ABT-199 or ABT-263. Annexin V^+^ cells were analysed by flow cytometry. Data are shown as mean ± s.d. (n = 3). Statistical significance of ABT-199 vs ABT-263 response was established by unpaired t-test for two group comparisons and defined as ****p<0.0001, ***p<0.001, **p<0.01, ns;p≥0.01. **D.** Kaplan-Meier plot showing the percentage of leukemia-free survival from vehicle vs ABT-263-treated NSG-SGM3 mice transplanted with M7012 PDX cells, over time. Statistical significance was measured by the log-rank test (Mantel-Cox), n=5. Upward ticks correspond to days of treatment; a white arrow corresponds to peripheral blood analysis (day 39 post-transplantation). **E.** Quantification of mCD45^-^hCD56^+^ cells from peripheral blood 39 days post-transplantation, or at sacrifice (if under 39 days posttransplantation).

We assessed the sensitivity of cells isolated from patient-derived xenografts to BH3 mimetics. ABT-263 (EC_50_<500nM) was significantly more potent than ABT-199 (EC_50_>10μM) at inducing apoptosis in cells isolated from patient-derived CBFA2T3-GLIS2^+^ xenografts (PDX; patients M7012 and M7014)(**Fig. 5C**)^5,38^. We then evaluated the *in vivo* efficacy of ABT-263 using this AMKL PDX model. Treatment with ABT-263 significantly prolonged survival compared to vehicle, with mice showing a median survival of 42 days versus 36 days, respectively (**Fig. 5D**). Also, ABT-263 strikingly reduced tumor burden, indicated by a significant decrease in human blasts in the peripheral blood (**Fig. 5E**). Taken together, observations from primary murine hematopoietic cells and CG2-driven leukemias, including human patient samples, are consistent with a cellular state of apoptotic priming in CBFA2T3-GLIS2^+^ AMKL, while the increased sensitivity to navitoclax further encourage the notion of a dependency on both BCL-xL and BCL-2 for survival of these cells.

## Discussion

Pre-clinical models that reliably reflect the molecular and phenotypical features of specific AML subtypes are necessary for the identification and testing of novel therapeutic strategies^23^. In the present study, we generated mouse models of AMKL driven by CBFA2T3-GLIS2 that shared striking similarities to the corresponding human leukemia^40^. Importantly, unlike a previously described inducible transgenic CBFA2T3-GLIS2 mouse model displaying an AML-skewed profile^10^, the retrovirus-based models we generated gave rise to, with high penetrance, leukemia with a consistent CD41^+^ AMKL phenotype. Of note, GLIS2, alone or fused to CBFA2T3, mediated the expression of megakaryoblast-specific genes *in vitro* and *in vivo*. Furthermore, we provide evidence that GLIS2, which alone is unable to transform HSPC *in vivo*, acts as an oncogene when co-expressed with activated Nras to promote the development of spontaneous AMKL phenotypically similar to CBFA2T3-GLIS2 AMKL. This suggests that GLIS2 constitutes an important driver of leukemogenesis in AMKL. Indeed, we revealed that CBFA2T3-GLIS2 and GLIS2 modulate the BCL2 family proteins Bcl2, Puma1, and Bim, consistent with apoptotic priming. BH3 mimetic treatment of CBFA2T3-GLIS2-driven murine AMKL cells and PDX cells revealed a striking sensitivity towards the BCL-2/BCL-x_L_ dual inhibitor navitoclax, both *in vitro* and *in vivo*. These results suggest that navitoclax may efficiently treat AMKL pediatric patients harboring the CBFA2T3-GLIS2 gene fusion.

One exceptional feature of the CBFA2T3-GLIS2 fusion is the dominant role for GLIS2 in driving the megakaryocytic phenotype. The actual cooperation between GLIS2 and activated Nras to generate AMKL is reminiscent of the “Multistep pathogenesis of AML” model proposed by Kelly and Gilliland^56^. In this model, class I mutations such as activated Ras may facilitate the development of GLIS2^+^ AMKL by promoting survival, proliferation and competitiveness of transplanted HSPC^57^. On the other hand, the capacity of GLIS2 to modulate the expression of anti-apoptotic genes while promoting self-renewal in HSPC^5^ suggests it may act as a class II oncogene. Notably, all of the CG2-Nras leukemias consistently displayed a luciferase signal which increased over time, in accordance with the fact that oncogenic Nras increases short- and long-term competitiveness in transplantation experiments^58,59^. CBFA2T3-GLIS2, which was sufficient to engage a Ras-associated transcriptional signature, presumably acts as both a class I and II oncogene, bypassing tumor-suppressor pathways, and acquiring the potential to produce aggressive AMKL in a single hit. This idea is supported by the paucity of hotspot mutations cooccurring with CBFA2T3-GLIS2^6^, and by the early age at which children present symptoms of AMKL^42^. Importantly, Nras^G12D^ accelerated the development of, and reduced the overall survival associated with CBFA2T3-GLIS2-driven leukemias, without affecting the phenotype. This is consistent with previous reports showing that activated Nras cooperates with other fusion proteins in human AML^24,44^, or with loss of *Trp53* and *Nf1* tumor suppressor genes, in a similar _fashion_^60,61^.

Additionally, our results demonstrated that human AMKL cell lines and murine HSPC as well as mouse and human leukemic cells expressing CBFA2T3-GLIS2 exhibit increased expression of BCL-2 family proteins in a manner indicative of apoptotic priming^52,53^. We suggest that CBFA2T3-GLIS2-driven leukemic cells are particularly sensitive to navitoclax and less so to venetoclax. Indeed, despite previous observations generating considerable optimism, venetoclax has shown limited effectiveness when used alone, particularly against relapsed or refractory AML^62^. Navitoclax, a potent BH3 mimetic and dual inhibitor of BCL-2/BCL-x_L_, is a promising drug for the treatment of leukemias^55,63^; nonetheless, the side effects of this drug, including anemia and thrombocytopenia, reduce its therapeutic utility^64^. Based on the structure of navitoclax, the reverse-engineered venetoclax, which specifically targets BCL-2 while having low affinity for BCL-x_L_, demonstrated reduced toxicity towards platelets *in vivo* and is generally well tolerated in AML patients^54,65–68^. Based on previous reports, a protective role for BCL-2 in cells from the megakaryocytic lineage is somewhat unexpected^54,69,70^. Assuming that BCL-2 is acting as the primary anti-apoptotic protein in CBFA2T3-GLIS2^+^ AMKL, venetoclax should have been at least as potent as navitoclax in inducing AMKL cell death. However, venetoclax induced only a weak apoptotic response (less than 50%) in human cell lines and PDX cells, while navitoclax efficiently promoted cell death at lower concentrations. Consequently, it is tempting to speculate that, in CBFA2T3-GLIS2^+^ AMKL cells, both BCL-2 and BCL-x_L_ together may provide protection against apoptosis induced by BH3 mimetics. Accordingly, the absence of either BCL-x_L_ or BCL-2 alone does not affect megakaryoblast or platelet survival, suggesting that their anti-apoptotic function may be complementary^69,71–73^. As such, AMKL cells expressing CBFA2T3-GLIS2 may be more sensitive to navitoclax because of i) their increased BCL-2 expression and ii) the inherent dependency of megakaryoblasts on BCL-x_L_ to prevent apoptosis. The prominent role of BCL-x_L_ in normal megakaryocyte biology combined with our data suggests navitoclax may serve as a superior BH3 mimetic in AMKL patients.

Overall, comprehensive immunophenotypic, transcriptomic, and molecular analysis of our CG2-driven mouse models of AMKL provided novel insights into the critical contribution of GLIS2 as a partner in the CBFA2T3-GLIS2 oncogenic fusion, and that the cells expressing this fusion likely take advantage of BCL-2 and BCL-x_L_ for protection against apoptosis. Our results support clinical consideration of navitoclax in the treatment of pediatric patients with CBFA2T3-GLIS2^+^ AMKL for which there is currently limited therapeutic options and a poor prognosis associated with this disease.

## Supporting information

Supplemental Material (3 Supp. figures + 3 Supp. Tables + Supp. Ref)

## Acknowledgements

The authors would like to thank Barry Cole and members of the Cole Foundation for their continuous support and for their efforts in promoting research in the fields of pediatric and young adult leukemia and lymphomas in Montréal, Qc (Canada). This work was supported by transition grants from the Cole Foundation (F.A.M., H.J.M., J.-S.D., F.E.M., L.H.), and operating grants from the Canadian Institutes of Health Research (PJT-156133; F.A.M., H.J.M., J.-S.D., E.A.D.), the Canadian Cancer Society-Quebec Division/Mont Gabriel Summit Research Fund/Cole Foundation (#705480; F.A.M., H.J.M., J.-S.D., E.A.D.), the Fondation de l’Hôpital Maisonneuve-Rosemont / Défi Vélo 2016 (F.A.M. and E.A.D.), and the Cancer Research Society (#25350; F.A.M. and L.H.). F.A.M. holds the Canada Research Chair in Epigenetics of Aging and Cancer. H.J.M., L.H. and J.S.D. hold career awards from the Fonds de Recherche du Québec - Santé (FRQS). M.N. obtained post-doctoral fellowships from the FRQS and the Cole Foundation. C-É.L-G., C.C., and K.B. were supported by post-doctoral fellowships from the Cole Foundation. C.S. obtained PhD studentships from the Cole Foundation, FRQS and Hydro-Québec.

## Authors’ contributions

**Conceptualization**: M. Neault, C. Capdevielle, C.É. Lebert-Ghali, H.J. Melichar and F.A. Mallette

**Data curation**: M. Neault, C.É. Lebert-Ghali, M. Fournier, H.J. Melichar and F.A. Mallette **Formal Analysis**: M. Neault, C.É. Lebert-Ghali, C. Capdevielle, M. Fournier, A. Obermayer, C. Sawchyn, J.S. Delisle, T.I. Shaw, T.A. Gruber, H.J. Melichar and F.A. Mallette

**Funding acquisition**: J.S. Delisle, E.A. Drobetsky, L. Hulea, H.J. Melichar and F.A. Mallette **Investigation**: M. Neault, C.É. Lebert-Ghali, C. Capdevielle, M. Fournier, E.A.R. Garfinkle, K. Nguyen, B. Assaf, T.A. Gruber, H.J. Melichar and F.A. Mallette

**Methodology**: M. Neault, C.É. Lebert-Ghali, C. Capdevielle, M. Fournier, A. Cotton, K. Boulay, F.E. Mercier, J.S. Delisle, E.A. Drobetsky, L. Hulea, J. Zuber, T.A. Gruber, H.J. Melichar and F.A. Mallette

**Project administration**: H.J. Melichar and F.A. Mallette

**Resources**: T.I. Shaw, T.A. Gruber, H.J. Melichar and F.A. Mallette

**Software**: M. Fournier, A. Obermayer, C. Sawchyn and T.I. Shaw

**Supervision**: E.A. Drobetsky, H.J. Melichar and F.A. Mallette

**Validation**: M. Neault, C.É. Lebert-Ghali, C. Capdevielle, M. Fournier, L. Hulea, H.J. Melichar and F.A. Mallette

**Visualization**: M. Neault, C.É. Lebert-Ghali, C. Capdevielle, M. Fournier, A. Obermayer **Writing – original draft**: M. Neault, H.J. Melichar and F.A. Mallette

**Writing – review & editing**: M. Neault, C.É. Lebert-Ghali, C. Capdevielle, M. Fournier, K. Boulay, J.S. Delisle, E.A. Drobetsky, L. Hulea, T.I. Shaw, J. Zuber, T.A. Gruber, H.J. Melichar and F.A. Mallette

## Notes

**Conflicts of interest:** The authors have declared that no conflict of interest exists.

### Competing Interest Statement

The authors have declared no competing interest.

