## Supplemental Material (3 Supp. figures + 3 Supp. Tables + Supp. Ref) for "CBFA2T3-GLIS2-dependent pediatric acute megakaryoblastic leukemia is driven by GLIS2 and sensitive to Navitoclax"

[1]Immunology-Oncology Unit, Maisonneuve-Rosemont Hospital Research Centre, Montréal, QC, Canada.; [2]Département de Microbiologie, Infectiologie et Immunologie, Université de Montréal, Montréal, QC, Canada; [3]Département de Biochimie et Médecine Moléculaire, Université de Montréal, Montréal, QC, Canada; [4]Department of Pediatrics, Division of Hematology, Oncology, Stem Cell Transplantation and Regenerative Medicine, Stanford University School of Medicine, Palo Alto, CA, USA; [5]Department of Biostatistics and Bioinformatics, H. Lee Moffitt Cancer Center, Tampa, FL, USA; [6]St. Jude Children's Research Hospital, Memphis, TN, USA; [7]Department of Medicine, McGill University, Montréal, QC, Canada; [8]Département de Médecine, Université de Montréal, Montréal, QC, Canada; [9]Research Institute of Molecular Pathology, Vienna, Austria.

**<u>Supplemental Material:</u> 3 Supplemental figures + 3 Supplemental tables + Supplemental References**

SUPPLEMENTAL FIGURE 1.

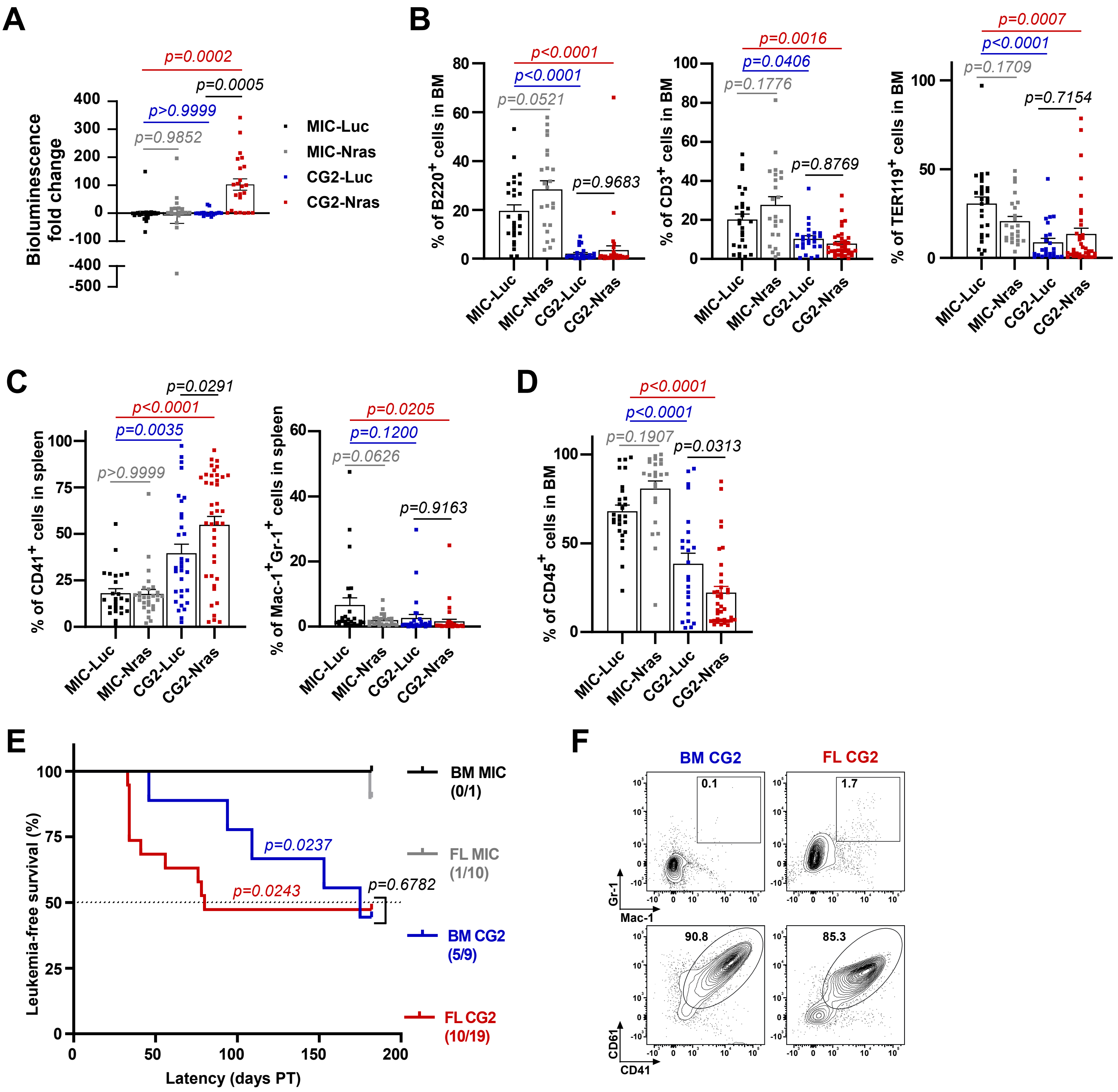

**Supplemental Figure 1. Characterization of CG2-Driven Leukemias.**
**A.** Relative fold change of bioluminescence (radiance) between first and last bioimaging, post-transplantation. **B, C.** Quantification of flow cytometry markers from mCherry$^+$ BM (**B**) and spleen (**C**) cells at time of sacrifice. **D.** Quantification of CD45$^+$ BM cells at time of sacrifice. **E.** Kaplan-Meier plot showing the percentage of leukemia-free survival from each vector group over time. Numbers represent leukemic mice over the total number of mice in each group. Statistical significance was measured by the log-rank test (Mantel-Cox) using FL MIC as a control. **F.** Representative flow cytometry plots of flow cytometry markers from mCherry$^+$ BM cells at time of sacrifice.

SUPPLEMENTAL FIGURE 2.

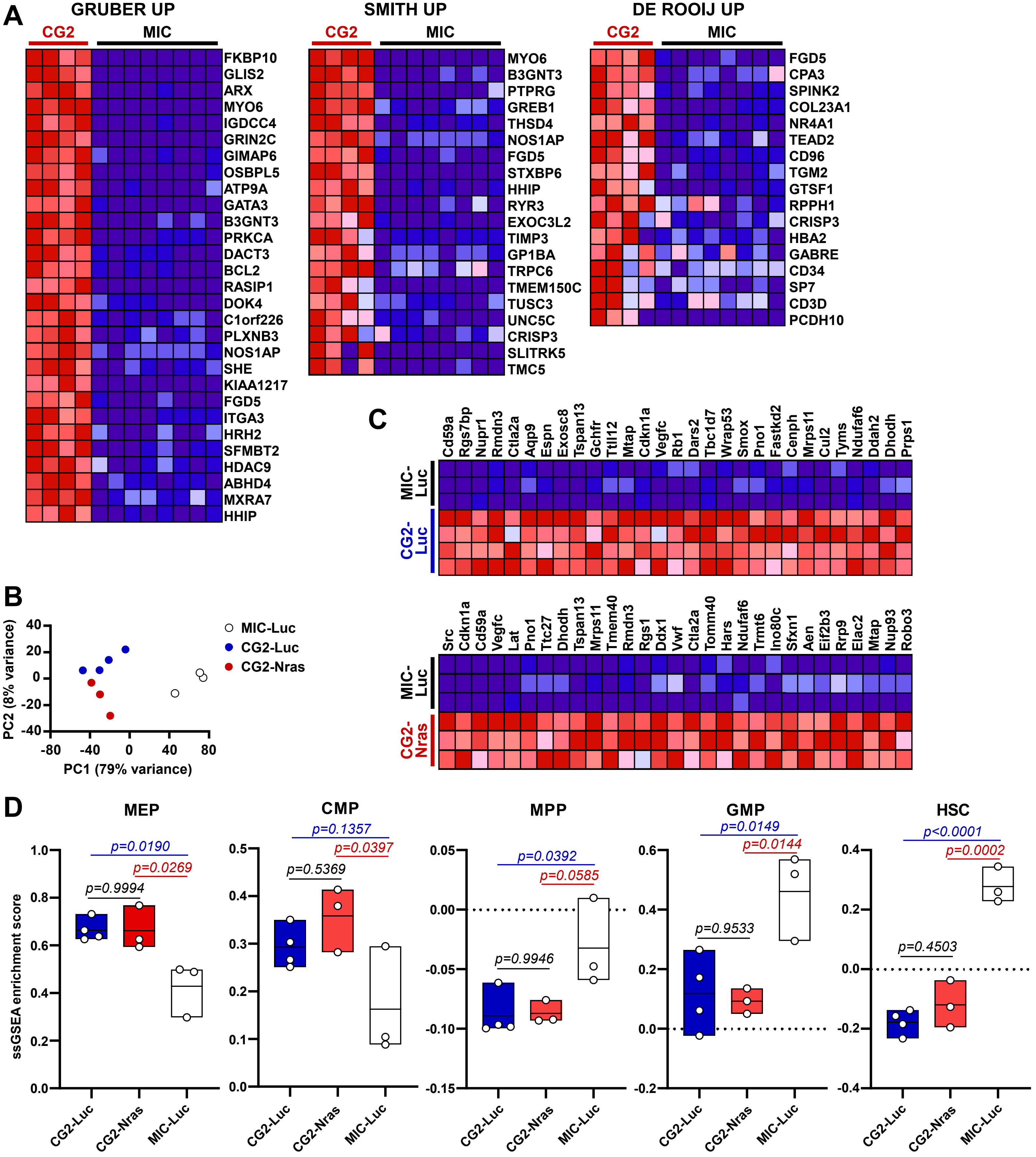

**Supplemental Figure 2. Pathway Enrichment and Myeloid Lineage Signature of CG2-Driven Leukemias.**

**A.** Heatmaps of the core enriched genes from FL cells expressing CBFA2T3-GLIS2 (CG2) vs control (MIC) for gene sets comprising genes upregulated in CBFA2T3-GLIS2$^{+}$ patients vs fusion-negative AMKL [1,2] or AML [3]. Increased expression (red), decreased expression (blue). The list of core enriched genes for the GRUBER UP gene set was truncated for better viewing (full list of genes is available in **Supplemental Table 2**). **B.** Principal component analysis (PCA) of the gene expression profiles from RNA-seq data described in **Fig. 2B**. **C.** Heatmap of the core enriched genes from mCherry$^{+}$-sorted BM cells of CG2-Luc and CG2-Nras at sacrifice, using MIC-Luc as a control for gene sets containing genes upregulated upon doxycycline induced expression CBFA2T3-GLIS2 FL HSPC vs WT HSC [4]. Increased expression (red), decreased expression (blue). The lists of core enriched genes were truncated for better viewing (full list of genes is available in **Supplemental Table 2**). **D.** Boxplots of myeloid lineage signature scores from **Fig. 2F**.

SUPPLEMENTAL FIGURE 3.

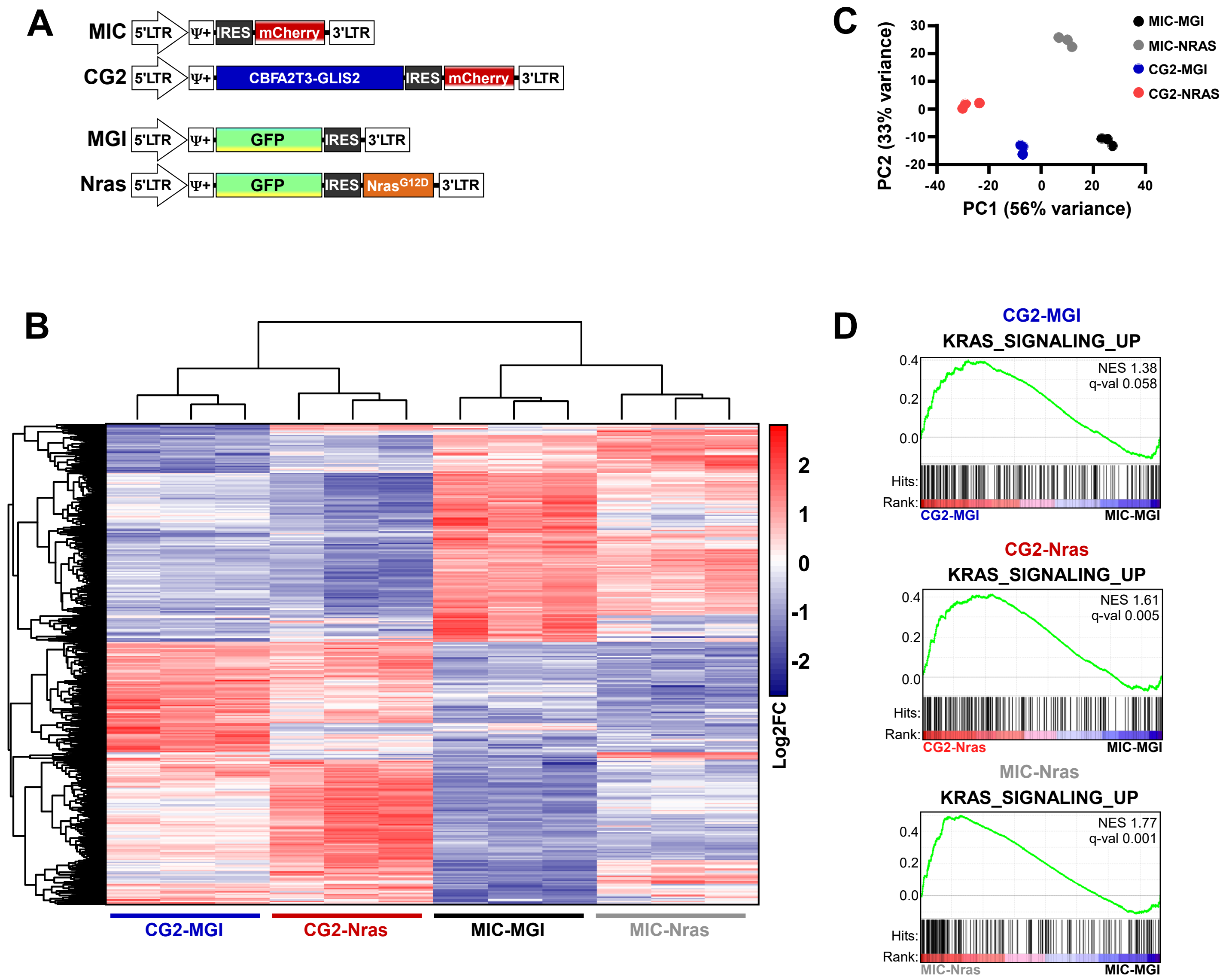

**Supplemental Figure 3. Ras Pathway Activation in CG2-Expressing HSPC.**
**A.** MSCV-based retroviral vectors were designed to be co-transduced in mouse FL cells. The mCherry fluorescent protein expresses CBFA2T3-GLIS2 or not, and the GFP construct expresses $Nras^{G12D}$ or not. FL cells were co-transduced with a combination of mCherry and GFP constructs. 48h post-transduction, FLcells were sorted for $mCherry^{+}GFP^{+}c\text{-}Kit^{+}lin^{-}$ (lin defined as $Gr1^{+}B220^{+}CD3^{+}TER\text{-}119^{+}$) expression and subjected to RNA-seq. **B, C.** Unsupervised hierarchical clustered heatmap of differentially expressed genes (most significant *p*-values; log2 fold change) (**B**) and PCA of the gene expression profiles (**C**) from sorted FL cells described in (**B**) for each vector combination. MIC-MGI was used as a control. **D.** GSEA plot of expression data obtained from sorted FL cells described in (**B**) showing enrichment of genes that are upregulated upon expression of oncogenic Ras [5].

**Supplementary Table 1. List of mouse strains, reagents and qPCR primers**

| Identifier | Reference # | Provider |
|---|---|---|
| ***Mice*** | | |
| C57BL/6J | 000664 | The Jackson Laboratory |
| B6.SJL-*Ptprc* $^{a}$ *Pepc* $^{b}$ /Boy$^{J}$ | 002014 | The Jackson Laboratory |
| NOD.Cg-Prkdc$^{scid}$ Il2r$^{tm1Wjl}$ Tg(CMV-IL2,CSF2,KITLG)1Eav/MloySzJ | 013062 | The Jackson Laboratory |
| ***Retroviral and lentiviral constructs*** | | |
| pMSCV-IRES-GFP (MIG) | 9044 | Addgene |
| pMSCV-Luciferase-IRES-Nras$^{G12D}$ (Nras) | 60834 | Addgene |
| pCL-ECO | 12371 | Addgene |
| pCW57.1 | 41393 | Addgene |
| pLentiCRISPRv2GFP | 82416 | Addgene |
| pMD2.G | 12259 | Addgene |
| psPAX2 | 12260 | Addgene |
| pENTR/D-TOPO | K2400-20 | Invitrogen |
| ***Flow cytometry antibodies and dyes*** | | |
| APC-anti-mCD117(c-kit) | 105812 | BioLegend |
| BV711-anti-mCD11b (Mac-1) | 101242 | BioLegend |
| APC/Cy7-anti-mLy-6G/Ly-6C(Gr1) | 108424 | BioLegend |
| BV605-anti-mCD41 | 133921 | BioLegend |
| PE/Cy7-anti-mCD61 | 104318 | BioLegend |
| APC/Cy7-anti-mCD45R/B220 | 103224 | BioLegend |
| PerCP/Cy5.5-anti-mCD3e | 100328 | BioLegend |
| BV785-anti-mTer119 | 116212 | BioLegend |
| Pacific Blue-anti-mCD45.1 (Ly-5.1) | 110722 | BioLegend |
| Alexa Fluor 700-anti-mCD45.2 (Ly-5.2) | 109822 | BioLegend |
| Biotin-anti-mLy-6G/Ly-6C(Gr1) | 108404 | BioLegend |
| Biotin-anti-mCD45R/B220 | 103204 | BioLegend |
| Biotin-anti-mCD3e | 100304 | BioLegend |
| Biotin-anti-mTER119 | 116204 | BioLegend |
| BV421-Streptavidin | 405225 | BioLegend |
| Pacific Blue-anti-mCD45 | 103126 | BioLegend |
| BV711-anti-hCD56 (NCAM) | 318335 | BioLegend |
| Zombi-AQUA™ | 423101 | Biolegend |
| APC-Annexin V | 640920 | Biolegend |
| ***Western blot antibodies*** | | |
| anti-CBFA2T3 | ab33072 | Abcam |
| anti-GLIS2 | PA5-40314 | Thermo Fisher Scientific |
| anti-Bcl2 | ab182858 | Abcam |
| anti-MCL1 | 5453 | Cell Signaling Technology |
| anti-BCL-$x_L$ | 2764 | Cell Signaling Technology |
| anti-BIM | ab32158 | Abcam |

| | | |
|---|---|---|
| anti-α-tubulin | T6074 | Sigma-Aldrich |
| anti-pan-actin | MS-1295-P0 | Thermo Fisher Scientific |
| HRP-Horse anti-mouse | 7076 | Cell Signaling Technology |
| HRP-Goat anti-rabbit | 7074 | Cell Signaling Technology |

***Real-time quantitative PCR (RT-qPCR) primers***

| Identifier | Sequence (5'-3') |
|---|---|
| ACTB-Fwd | TCAAGATCATTGCTCCTCCTGAG |
| ACTB-Rev | ACATCTGCTGGAAGGTGGAC |
| PPIA-Fwd | GTCAACCCCACCGTGTTCTT |
| PPIA-Rev | CTGCTGTCTTTGGGACCTTGT |
| CBFA2T3-Fwd | AGAAAGCCGTGTCGGACG |
| GLIS2-Rev | AAATAGCGCAGTGGCTGGAAG |
| BBC3-Fwd | CGAGATGGAGCCCAATTAGGTG |
| BBC3-Rev | TACATGGTGCAGAGAAAGTCCC |
| BCL2a-Fwd | TGATTTCTCCTGGCTGTCTCTG |
| BCL2a-Rev | TTGCATATTTGTTTGGGGCAGG |
| BCL2L1-Fwd | GGCCACTTACCTGAATGACC |
| BCL2L1-Rev | AAGAGTGAGCCCAGCAGAAC |
| BCL2L11-Fwd | GCATCATCGCGGTATTCGGT |
| BCL2L11-Rev | CATCAGAAGGTTGCTTTGCCAT |
| BMP2-Fwd | GCTAGACCTGTATCGCAGGC |
| BMP2-Rev | CCAAAGATTCTTCATGGTGGAAGC |
| ERG-Fwd | GCTGCTCAACCATCTCCTTC |
| ERG-Rev | ACAGGAGCTCCAGGAGGAAC |
| GATA1-Fwd | GACAGGCCACTACCTATGCAA |
| GATA1-Rev | TGCCCGTTTACTGACAATCAGG |
| ID1-Fwd | GCCCCAGAACCGCAAGGTGA |
| ID1-Rev | AGGAACGCATGCCGCCTCG |
| MCL1-Fwd | TCTCTCGGTACCTTCGGGAG |
| MCL1-Rev | TTGATGTCCAGTTTCCGAAGCAT |

### Supplementary Table 2. List of GSEA core enriched genes

| GRUBER UP | SMITH UP | DE ROOIJ UP | LOPEZ UP | | ROSS AML Of FAB M7 TYPE | |
|---|---|---|---|---|---|---|
| FL CG2-Luc vs FL MIC-Luc | | | BM CG2-Luc vs BM MIC-Luc | BM CG2-Nras vs BM MIC-Luc | BM CG2-Luc vs BM MIC-Luc | BM CG2-Nras vs BM MIC-Luc |
| FKBP10 | MYO6 | FGD5 | Cd59a | Src | GP1BA | LAT |
| GLIS2 | B3GNT3 | CPA3 | Rgs7bp | Cdkn1a | BMP2K | BMP2K |
| ARX | PTPRG | SPINK2 | Nupr1 | Cd59a | MYH10 | GP1BA |
| MYO6 | GREB1 | COL23A1 | Rmdn3 | Vegfc | GP1BB | SOD1 |
| IGDCC4 | THSD4 | NR4A1 | Ctla2a | Lat | SLC39A4 | TNIK |
| GRIN2C | NOS1AP | TEAD2 | Aqp9 | Pno1 | GATA1 | HIGD1A |
| GIMAP6 | FGD5 | CD96 | Espn | Ttc27 | TNIK | GATA1 |
| OSBPL5 | STXBP6 | TGM2 | Exosc8 | Dhodh | SOD1 | TPM1 |
| ATP9A | HHIP | GTSF1 | Tspan13 | Tspan13 | APOE | GP1BB |
| GATA3 | RYR3 | RPPH1 | Gchfr | Mrps11 | DNAJC6 | PROS1 |
| B3GNT3 | EXOC3L2 | CRISP3 | Ttll12 | Tmem40 | MINPP1 | PCCB |
| PRKCA | TIMP3 | HBA2 | Mtap | Rmdn3 | TPM1 | SLC39A4 |
| DACT3 | GP1BA | GABRE | Cdkn1a | Rgs1 | CMAS | ATP5MPL |
| BCL2 | TRPC6 | CD34 | Vegfc | Ddx1 | NDUFB6 | NDUFB6 |
| RASIP1 | TMEM150C | SP7 | Rb1 | Vwf | TAL1 | GATA2 |
| DOK4 | TUSC3 | CD3D | Dars2 | Ctla2a | RDX | PRUNE1 |
| C1orf226 | UNC5C | | Tbc1d7 | Tomm40 | HIGD1A | MRPS12 |
| PLXNB3 | CRISP3 | | Wrap53 | Hars | ICAM4 | TAL1 |
| NOS1AP | SLITRK5 | | Smox | Ndufaf6 | ATP5MPL | CUTA |
| SHE | TMC5 | | Pno1 | Trmt6 | DRAP1 | ITGA2B |
| KIAA1217 | | | Fastkd2 | Ino80c | MRPS12 | BEX3 |
| FGD5 | | | Cenph | Sfxn1 | ITGA2B | MYH10 |
| ITGA3 | | | Mrps11 | Aen | PLOD2 | DNAJC6 |
| HRH2 | | | Cul2 | Eif2b3 | ABCC4 | SERPINI1 |
| SFMBT2 | | | Tyms | Rrp9 | CUTA | PLOD2 |
| HDAC9 | | | Ndufaf6 | Elac2 | ZMYND8 | ABCC4 |
| ABHD4 | | | Ddah2 | Mtap | PROS1 | APOE |
| MXRA7 | | | Dhodh | Nup93 | TFR2 | CMAS |
| HHIP | | | Prps1 | Robo3 | TIMP3 | PSMD6 |
| ALS2CL | | | Ddx1 | Lyar | PRUNE1 | MINPP1 |
| ANO8 | | | Akr1c13 | Derl1 | | |
| FRMD4B | | | Taf9 | Mphosph6 | | |
| ECM1 | | | Xpnpep1 | Cdc34 | | |
| LTC4S | | | Akip1 | Espn | | |
| PLXNB1 | | | Ino80c | Tex30 | | |
| FAM171A2 | | | Apoe | Dars2 | | |
| TNFSF11 | | | Dscc1 | Idh3a | | |
| ACR | | | Hars | Rpf2 | | |
| ENPP6 | | | Nbn | Ctps | | |
| CDH5 | | | Agpat5 | Haus7 | | |
| LPAR6 | | | Pdss1 | Prmt7 | | |
| QRFPR | | | Alg8 | Nbn | | |
| CDH2 | | | Ttc27 | Ppp2r1b | | |
| MAGEE1 | | | Ptpn11 | Noc4l | | |
| C7orf57 | | | Eif5al3-ps | Lars | | |
| BIK | | | Hprt | Taf9 | | |
| KIF1C | | | Nup205 | Sapcd2 | | |
| PCDH19 | | | Txnrd2 | Abce1 | | |
| HSD17B14 | | | 2410004B18Rik | Mrps18b | | |
| PHLDB1 | | | Siah1b | Ndufaf1 | | |
| ZDBF2 | | | Eif2a | Gspt1 | | |
| ZNF608 | | | Mphosph6 | Ddx18 | | |
| DPYSL2 | | | Prmt7 | Mrto4 | | |
| EXOC3L2 | | | Chaf1b | Hacd3 | | |
| SPINK2 | | | Sfxn1 | Gfm1 | | |
| ST6GALNAC3 | | | Wdr36 | Wdr75 | | |
| AIF1L | | | Coa5 | Exosc9 | | |
| SLC7A8 | | | Aifm1 | Ftsj3 | | |
| CDH24 | | | Trmt6 | Nudt1 | | |
| SPOCK2 | | | Cdk6 | Rrp12 | | |
| TEAD2 | | | Tomm40 | Eif2b1 | | |
| TNFRSF11A | | | Rfc5 | Wdr36 | | |
| AMIGO1 | | | Mars | Pop1 | | |
| SPAG1 | | | Minpp1 | Wrap53 | | |
| GPC2 | | | B3galt6 | Nif3l1 | | |
| STOX2 | | | Lmna | Tbl3 | | |

| | | |
|---|---|---|
| CNIH3 | Csf2rb2 | Pmpca |
| IL1R2 | Lars | Ptcd3 |
| SCN1B | Cluh | Cul2 |
| MKRN3 | Atic | Cad |
| BMP4 | Tex30 | Alg8 |
| TMEM25 | Dis3 | Trmt11 |
| TNXB | Gspt1 | Fastkd2 |
| GSN | Cenpn | Nle1 |
| SPTAN1 | Exosc9 | Tuba8 |
| HPN | Ptcd3 | Thop1 |
| IGFBP7 | Gmps | Psmg3 |
| APLN | Elac2 | Pdss1 |
| TNFAIP3 | Ctps | Tars2 |
| FZD8 | Derl1 | Steap3 |
| SRGAP3 | Nup85 | Ttll12 |
| ZNF704 | Idh3a | Pla2g4a |
| PRX | Rcl1 | Gmps |
| TPM4 | Pmpca | Xpnpep1 |
| MMP2 | Clpb | Fermt3 |
| PLCE1 | Polr1e | Cenph |
| PTPRB | Ndufs1 | Spryd4 |
| CHSY1 | Ddx18 | Nsun2 |
| DUSP27 | Nup93 | Tyms |
| STAP2 | Nsun2 | 2410004B18Rik |
| PDZD2 | Mcm8 | Gchfr |
| RTN4R | Tk1 | Gtse1 |
| CXXC4 | Nup43 | Rfc5 |
| ABCA5 | Nans | Parn |
| EPHB2 | Tmem97 | Nolc1 |
| MMP14 | Rabggtb | Cyb561d2 |
| MRAS | Nudt1 | Nop9 |
| PTGER1 | Thg1l | Zpr1 |
| LMTK3 | Ap3s2 | Polr1e |
| RAB11FIP4 | Fignl1 | Snrpa1 |
| ACVRL1 | Ubox5 | Cluh |
| TUSC3 | Parn | Sars2 |
| SALL2 | Rad18 | Tbc1d7 |
| C1orf115 | Plaa | Bysl |
| TMEM150B | Ankrd27 | Thg1l |
| PRR5L | Haus7 | Gnl2 |
| UNC5C | Mrto4 | Rcl1 |
| CRISP3 | Mre11a | Atic |
| PLXNB2 | Gfm1 | Coa5 |
| CD40 | Wdr75 | Mrpl22 |
| EPHB6 | Trmt11 | Rpap3 |
| CMTM3 | Utp18 | Akip1 |
| NDST3 | Tdp1 | Plaa |
| CDH13 | Suv39h1 | Csf2rb2 |
| THSD1 | Tars2 | Supv3l1 |
| MORC4 | Tcrg-C1 | Eif5al3-ps |
| DUSP10 | Mecr | Lmna |
| VDR | Pnpt1 | Naa25 |
| MAPK11 | Mipep | Heatr3 |
| MPPED2 | Lyar | Nans |
| MYO1B | Mthfd1 | Gemin6 |
| ADAMTS17 | Nup107 | Aifm1 |
| CPNE2 | Dph5 | Ormdl2 |
| SH3RF3 | Sars2 | Tdp1 |
| SPATA18 | Robo3 | Nup85 |
| PNPLA3 | Thoc1 | Ankrd27 |
| PCDH12 | Opa1 | Exosc8 |
| NUAK1 | Blvra | Ndufs1 |
| SPNS2 | Cenpm | Yrdc |
| TPD52L1 | Chd1l | Nom1 |
| ST8SIA2 | Tmx2 | Tbrg4 |
| LPIN3 | Zfp930 | Nol11 |
| SPTBN5 | Ppat | Ptpn7 |
| EFNA1 | Rpp30 | Mcm8 |
| PCSK5 | Ndufaf1 | Etf1 |

| | | |
|---|---|---|
| PPP1R13B | Oip5 | Ddx56 |
| CD34 | Ttk | Dcaf13 |
| ILDR2 | G6pc3 | Rad18 |
| WT1 | Rrp9 | Rcc1l |
| UBE2E2 | Orc3 | Dlat |
| KITLG | Supv3l1 | Utp18 |
| SP7 | Zpr1 | Mars |
| CD3D | Mrps18b | Ptpn11 |
| PTPRM | Mettl1 | Hspbp1 |
| RGS3 | Abce1 | Ubox5 |
| SOX11 | Dcaf13 | Prune1 |
| ERRFI1 | Cad | Dis3 |
| PCDH10 | Aen | Pes1 |
| COL6A3 | Pom121 | Ecd |
| ITM2A | Il15 | Fam98a |
| CCL1 | Tamm41 | Rb1 |
| PGF | Src | Lsg1 |
| UBE2QL1 | Hlf | Dph5 |
| RHOJ | Ftsj3 | Mrps30 |
| ABCA2 | Orc2 | Nat10 |
| SMAD9 | Mrps30 | Hacd1 |
| | Ppp2r1b | Thoc1 |
| | Ahcy | Surf2 |
| | Nom1 | Hprt |
| | Mrps2 | Ddx21 |
| | Rpf2 | Prim2 |
| | Mrpl22 | Mmp14 |
| | Polr3g | Nup37 |
| | Etf1 | Ap3s2 |
| | Eif2b3 | Mre11a |
| | Ska3 | Ipo4 |
| | Thop1 | Prps1 |
| | Steap3 | Tamm41 |
| | Gins1 | Mrpl35 |
| | Nomo1 | Atad3a |
| | Pla2g4a | Igf2bp3 |
| | Fen1 | Wdr55 |
| | Pusl1 | Gtf2f2 |
| | Mmp14 | Thoc6 |
| | Hells | Gtf2h1 |
| | Hdhd2 | Alox5 |
| | Pde12 | Nup205 |
| | Tubg1 | Nup43 |
| | Nol11 | Champ1 |
| | Sparc | Ahcy |
| | Hacd3 | Galk1 |
| | Trip13 | Wdr4 |
| | Anapc15 | Nup107 |
| | Coq7 | Dnajc11 |
| | Thap12 | Ddah2 |
| | Prim2 | Tubg1 |
| | Med17 | Chchd4 |
| | Gnl2 | Rars |
| | Alg3 | Qars |
| | Nop9 | Chd1l |
| | Ormdl2 | Mogs |
| | Tmem40 | Gart |
| | Snrpa1 | Mettl1 |
| | Alox5 | Cenpn |
| | Echdc2 | Pja1 |
| | Naa25 | Ppat |
| | Kcnn4 | Qtrt1 |
| | Rad51 | Bop1 |
| | Ecd | Hdhd2 |
| | Bysl | Eif2b5 |
| | Spdl1 | P2rx1 |
| | Me2 | Suv39h1 |
| | Ccnb1 | Pusl1 |
| | Srm | Alg3 |

| | |
|---|---|
| Dimt1 | Mrpl46 |
| Art4 | Eif2a |
| Rcc1l | Ddx27 |
| Heatr3 | Mrps2 |
| Ddx39 | Polr1b |
| Nle1 | Vldlr |
| Utp6 | Pmm2 |
| Atad3a | Dimt1 |
| Gtse1 | Mrps27 |
| Idi1 | Heatr1 |
| Heatr1 | Blvra |
| Galk1 | Polr3d |
| Ndc1 | Utp4 |
| Wdr46 | Usp10 |
| Nt5c3 | Rabggtb |
| Plscr2 | Rrp8 |
| Esco2 | Coq7 |
| Rrp12 | Med17 |
| Mrpl35 | Orc2 |
| Wdr55 | Pde12 |
| Exo1 | Ints5 |
| Dlat | Wdr46 |
| Dnajc11 | Aatf |
| Cdc34 | Anapc15 |
| Lonp1 | Orc3 |
| Ppih | Lmo2 |
| Mrpl46 | Pgam5 |
| Get4 | Mlkl |
| Sephs2 | F2rl3 |
| Hspbp1 | Rpp30 |
| Tbl3 | Lonp1 |
| Aunip | B3galt6 |
| Noc4l | Prpf38a |
| Mad2l1 | Gpd2 |
| Ddx21 | Slc25a10 |
| Wdr4 | Cdk6 |
| Bms1 | Tcrg-C1 |
| Nat10 | Tsr1 |
| Sapcd2 | Grwd1 |
| Cyb561d2 | Npm3 |
| Pes1 | Nol9 |
| Ddx56 | Mrpl16 |
| Rfc4 | Opa1 |
| Nolc1 | Trmt61a |
| Tbrg4 | Mecr |
| Phka2 | Mipep |
| Slc43a3 | Nop2 |
| Gmnn | Nt5c3 |
| Bub1 | Apoe |
| Gemin6 | Timm44 |
| Eif2b1 | Elp3 |
| Mrps10 | Gpatch4 |
| Fxr1 | Rrp15 |
| Mlkl | Sephs2 |
| Cdc27 | Smox |
| Nif3l1 | Agpat5 |
| Dcps | Pnpt1 |
| Thoc6 | Ppan |
| Exosc10 | Pdcd11 |
| Prpf38a | Bms1 |
| Mthfd2 | Rrp1b |
| Mrps27 | Fam136a |
| Spns3 | Fpgs |
| Rrp1b | Chaf1b |
| Psmg3 | Pex16 |
| Utp15 | Ddx39 |
| Hist1h2ae | Ecsit |
| Suv39h2 | Trip13 |
| Ckap2 | Ccdc22 |

Rars
Mrpl16
Mki67
Npm3
Tsr1
Riox2
Mns1
Fpgs
Acad9
Nup133
Vwf
Pmm2
Spryd4
Prune1
Pop1
Pus10
Kpna3
Usp10
Timm44
Nol9
Hmmr
Polr1b
Cdc6
Kif20b
Ddx27
Knl1
Mmachc
Rtel1
Stard4
Cdca2
Gart
Gm4737
Cdc25c
Rtca
Shcbp1
Cenpa
Nefh
Kpna2
Mrps31
Gtf2f2
Lrwd1
Mbd3
Cdk1
Rangrf
Nploc4
Pex16
Qars
Dhrs13
Mcm5
Rpap3
Gpatch4
Sppl2b
Mis18bp1
Rad54l
Elp3

Nploc4
L3mbtl2
Katnb1
Mbd3
Minpp1
Get4
Polr3g
Srm
Emilin2
Mthfd1
Rtca
Zfp930
Faap24
Asna1
Nomo1
Las1l
Rbm19
Tmx2
Mrps7
Sparc
Dkkl1
Utp6
Mthfd2
Siah1b
Utp15
Mns1
Nol12
Unc13d
Ppp1r8
Dscc1
Mrps18a
Mrpl20
Nefh
Clpb
Mrpl9
Trmt1
Fignl1
Mmp2
Srp54b
Farsa
Fxr1
Mphosph10
Enoph1
Id2
Aqr
Mrps10
Lrrc59
Mmachc
Mad2l1
Nup133
Hist1h4i
Exosc10
Nubp2
Nubp1
Traip
Dcps
Ccdc25
Mettl16
Kpna3
Ccnb1
Rgs7bp
Pus10
Setd6
Senp3
Ddx49
Utp14a
Tmem97
Babam1
Ndufaf4

Coq5
Lrwd1
Kcnn4
Ltv1
Prpf4
Nup35
Plscr2
Cox18
Cox10
Yars2
Trnt1
Odc1
Dok1
Wdr34
Bid
Rfc4
Dph3
Nup50
Rtel1
Mrps31
Taf11
Zfp64
Psmd13
Dhrs13
Ndc1
Acad9
Gtf2f1
Cinp
Timm22
Telo2
Supt16
Bcs1l
Ttk
Mmadhc
Rnf126
Tufm
Cdc27
Hlf
Tfam
Pom121
Il2rg
Suv39h2
Me2
Pwp2
Gss

### Supplementary Table 3. Myeloid Lineages Gene Sets

For the AML lineage analysis, myeloid lineage gene sets were derived from RNAseq samples of flow-sorted mouse hematopoietic progenitors, including HSC, GMP, CMP, MEP, and MPP as previously described in Dang et al. Leukemia 2017 (PMID: 28174417). LIMMA was used to define differentially expressed genes with adj.pval < 0.05. log2FC cutoffs were more stringent in HSC samples due to the high number of differentially expressed genes. Log2FC cutoff was lowered in CMP samples to capture more differentially expressed genes.

| HSC [Cutoff >2log2FC] | CMP [ Cutoff >0.5log2FC] | MPP [Cutoff >1log2FC] | GMP [Cutoff >1log2FC] | MEP [Cutoff >1log2FC] |
|---|---|---|---|---|
| Gm10391 | Kcng1 | Lrrn4 | Ear6 | Mc2r |
| Igkv4-58 | Angptl2 | Pcp4l1 | 1700012B09Rik | Rhag |
| Gm10394 | Vit | Igdcc4 | Ldhc | Sh3tc2 |
| Klhl14 | Pf4 | Nkx2-3 | Fam83a | Slc25a21 |
| Ighv2-9 | Homer2 | Edar | Ear1 | Tnfaip2 |
| Fcrl5 | F2rl2 | Msi2 | Prg3 | Nags |
| Scn4a | Itga2b | Hnf4a | Gm8113 | Ppm1l |
| Cd8b1 | Epha7 | Scube3 | Ear2 | Gclm |
| Gm8818 | Treml1 | 9030619P08Rik | Nrg1 | Cldn13 |
| Duxbl | Upk1b | Hlf | Prg2 | Abcg4 |
| Klk1 | Gucy1b3 | Epb4.1l4a | Plcb1 | Btnl10 |
| Gad1 | Bcl2 | Crispld1 | Tctex1d1 | Tmem56 |
| Faim3 | Myom1 | Il1r1 | Epx | Spire1 |
| Igkv13-84 | Rbpms2 | Il17re | Mfsd7a | Pklr |
| Cr2 | Lhfp | Sgsm1 | Matn2 | Epdr1 |
| Ighv1-19 | Pde3a | Kcnh2 | Hal | Trim2 |
| Igkv8-27 | Tuba8 | Bcam | Slc36a2 | Sphk1 |
| Cxcr5 | Flnb | Mecom | Depdc7 | Mapk9 |
| Scd1 | Mdm1 | Spint1 | Stxbp6 | A730046J19Rik |
| Dtx1 | Gp1bb | Cd276 | Slc30a2 | Sh2d4a |
| Igkv4-72 | Padi2 | Bdh2 | Tmem178 | Gfi1b |
| Mybpc2 | Gzmb | 1500009L16Rik | Mcpt8 | Cyth3 |
| A630023P12Rik | Sec31b | Gpr161 | Papss2 | Gm10430 |
| Ms4a1 | Ppfia4 | Myof | Cux2 | 1190007F08Rik |
| Ccr7 | Stbd1 | Cyp2j9 | Mcf2l | Sept8 |
| Ighv2-6 | Ifi205 | Ppp1r9a | Cyp11a1 | Atp1b2 |
| Il1f9 | | Dst | Trim45 | Myo1d |
| 4930426D05Rik | | Casp12 | F7 | Atp7b |
| Igkv10-95 | | Mmp2 | Mefv | Fam59a |
| Foxp3 | | Cand2 | Arhgef10l | 8430419L09Rik |
| Agpat9 | | Slc16a12 | F10 | Ermap |
| Fcamr | | Zfp532 | Gstm4 | Epb4.9 |
| Igkv1-88 | | Kcna2 | F13a1 | Tspan8 |
| Sit1 | | Thsd1 | Prss16 | Clstn1 |
| Cpm | | Slco2b1 | Lamp1 | Tal1 |
| CAAA01098150.1.2208.1 | | Laptm4b | Aldh1b1 | 1110020G09Rik |
| Ighv1-64 | | Mpl | 0610040J01Rik | C530008M17Rik |
| Ighg3 | | 1600021P15Rik | Lypd6 | Ptpn14 |
| Fam83g | | Epb4.1l1 | Alas1 | Pm20d2 |
| Gm8369 | | Samd12 | Sncaip | Aqp1 |
| Havcr1 | | Gas6 | Rnase12 | Slc26a1 |
| Tmem132e | | Vldlr | Fkbp1b | Foxh1 |
| Zbtb32 | | Mertk | Rnf43 | B3galtl |
| Il2ra | | Abcc3 | Trem2 | Add2 |
| Igkj1 | | Sema3d | Anxa3 | Stx2 |
| Ctla4 | | Gm16485 | Prss34 | Prss50 |

| | | | |
|---|---|---|---|
| Lta | Cyp2j6 | Svip | Fpgs |
| Igkv8-30 | Krt7 | Slc44a4 | Hemgn |
| Sspn | Siglec1 | Gpr160 | Ces2g |
| Cecr2 | Dennd2a | Met | Asph |
| Ighv5-9 | Paqr5 | Tmed3 | Acsl6 |
| Ighv9-3 | Ddx4 | Mmp19 | Fam126a |
| Ccr8 | Gm3739 | Tfec | Aldh1a7 |
| Cd19 | Emcn | Itga1 | Slc38a5 |
| Grm8 | Apob | Dmkn | Acmsd |
| Ighv5-16 | C77080 | Acpl2 | Spna1 |
| Igkv6-17 | Cyp7b1 | Ly6c1 | Igsf3 |
| Folr4 | Msrb3 | Soat1 | Car2 |
| Adm | Stxbp4 | Fcer1a | Fam132a |
| Scn4b | Hoxa9 | Idh1 | 1700063H04Rik |
| Cacna1i | Vnn1 | Ctsg | Lrrc8c |
| Sbk1 | Fam171a1 | B3gnt8 | Myh10 |
| Gm15326 | Axl | Bzrap1 | Ank1 |
| Lef1 | C1ra | Selm | Epb4.2 |
| Il9r | Parp12 | Apba1 | Ap1b1 |
| Cd209d | Alpk3 | Prss57 | Gls2 |
| Fcer2a | Sall2 | Grb14 | Minpp1 |
| Fam129c | BC067074 | Ly6c2 | Nipa1 |
| Gm10552 | Ttc8 | Gstm1 | Rhd |
| Ighv5-6 | Camsap3 | Agps | Zfpm1 |
| Prkcc | Col16a1 | Cd63 | Fam55b |
| Spib | Adam22 | Prtn3 | Ttll12 |
| Bcar3 | Basp1 | 4933440M02Rik | Agtr1a |
| Ighv1-78 | Zfp618 | Clec12a | Ntn4 |
| Ffar1 | Slc27a6 | Mgl2 | Mboat2 |
| Cacna1s | Gyltl1b | Cd63-ps | Ephx2 |
| Ltk | Pitx2 | Ptgr1 | Slc6a20a |
| Ighv9-4 | Hdac11 | Cldn15 | Shank3 |
| Bank1 | Csgalnact1 | Elane | Abcb4 |
| Irf4 | Sstr2 | Gpc1 | 3110056O03Rik |
| Cplx2 | Ifi44 | Afap1 | Parm1 |
| Cd40 | Glis2 | Ms4a3 | Col5a1 |
| 1810046K07Rik | Col14a1 | Chst13 | Optn |
| Vpreb3 | Cdcp1 | Med21 | Ugcg |
| Igkv8-28 | C030037D09Rik | Atp6v1b2 | Mylk3 |
| Chst3 | Gprasp2 | Fcgr3 | Arhgef25 |
| Mapk11 | Ccdc112 | Plekhh2 | Aldh1a1 |
| Cacna1e | Eya2 | Mmp28 | Sowaha |
| Gm16015 | Tgtp1 | C1galt1c1 | Pigq |
| Gm15408 | Epb4.1l3 | B4galt6 | Klf1 |
| Ighv5-4 | Gm4841 | Arsb | Mfsd2b |
| Igkv4-53 | Wbscr27 | Gm6713 | Acsl5 |
| Cd4 | Sel1l3 | Mgam | Car1 |
| Ighv1-5 | Cadps2 | | Scrn3 |
| Stac2 | BC068157 | | Cpox |
| Pax5 | Fstl1 | | Spon2 |
| Foxo1 | Dock3 | | 1300017J02Rik |
| Alpl | Tspan6 | | Kel |
| Icos | Itgb5 | | Asns |

| | | |
|---|---|---|
| Gm20506 | Hoxa10 | Sec14l2 |
| H2-Eb2 | E130304F04Rik | Trib2 |
| Igkv3-10 | Prdm16 | 9430076G02Rik |
| Ighv3-5 | Oasl2 | Gata1 |
| Cd79a | Itgad | Stxbp1 |
| Igkv1-117 | Postn | B3galnt2 |
| Cd8a | Rfx8 | 6030468B19Rik |
| Lynx1 | B4galnt4 | Hk1 |
| Crmp1 | Fam69b | Spnb1 |
| Ighv5-2 | Kcnj10 | Aldh18a1 |
| Myo1e | Gm13986 | Slc43a1 |
| Tnfrsf19 | Fcna | Cul4a |
| Nt5e | Gm16897 | Adcy6 |
| Dennd5b | Nlrp10 | Ptpn13 |
| 2210403K04Rik | Gcnt2 | Ammecr1 |
| Dll4 | Actn2 | Orc5 |
| Igkv8-19 | Tnip3 | Mt2 |
| Il10 | Ltbp3 | Acss2 |
| Igkv4-55 | Gm20467 | Acss1 |
| Ighv1-7 | 4930452B06Rik | Fam55d |
| Ighv8-12 | Tfcp2l1 | Pkhd1l1 |
| St6gal1 | Fam190a | Phyhip |
| Fam70a | Gm13881 | Aco1 |
| Igkj4 | Klhl4 | Samd14 |
| Cxcr7 | Maged2 | Plscr1 |
| Ighv10-1 | Epb4.1l4b | Tmem120b |
| Slc12a3 | Prkaa2 | Rab4a |
| Pou2af1 | Armcx1 | Hpn |
| Ighv1-84 | Hoxa3 | Gypa |
| Wnt10a | Nsg1 | Pla2g4c |
| Igkv1-135 | Nsun7 | Ctse |
| Tnfrsf13c | Vipr2 | Nfia |
| Pltp | Gm3696 | Steap3 |
| Ighv11-1 | Gpd1 | Grb10 |
| Col27a1 | Clic5 | Reep6 |
| Slamf6 | Serpinb2 | Spire2 |
| Trac | Sgce | 2200002K05Rik |
| Igkj2 | Ccdc48 | Zg16 |
| Stab2 | Rbpms | Lmcd1 |
| Ighv1-75 | Lrrc49 | Ltbp1 |
| Ighv5-9-1 | Ociad2 | Ccne1 |
| Gm16170 | Trpc6 | Pax9 |
| Ebf1 | Procr | Parvb |
| Vpreb1 | Ggt5 | Golm1 |
| Akap12 | Dlg4 | Slc41a3 |
| Ighv1-15 | Cdk18 | Ptdss2 |
| Pydc3 | Ldhd | Trim58 |
| Ighv8-5 | Vsig10 | Slc11a2 |
| Igkj3 | Gm5111 | Pcyt1b |
| Ighv6-3 | Alox15 | Slc22a23 |
| Dnase1l3 | Cd5l | Muted |
| Cacna1h | Ift81 | Clybl |
| Il1b | Nlrp1b | Ssx2ip |

| | | |
|---|---|---|
| Rgs16 | Camk2a | Xrcc5 |
| E330020D12Rik | Islr | Mmp14 |
| Igkv16-104 | Rims3 | Fgfrl1 |
| A530040E14Rik | Myo5c | Abcb6 |
| Igll1 | Slc16a11 | Sun1 |
| Ighv1-54 | Adc | Ifrd2 |
| Rasgrp1 | Ypel1 | Mns1 |
| Cdkl1 | Arhgef17 | Wdr60 |
| Ighv11-2 | Pkia | Nckap1 |
| Pacsin1 | Slc6a1 | Bex4 |
| Bcl11b | Calcrl | Tspan33 |
| Ighv9-1 | C1qc | Mthfd1 |
| Iglv3 | Il17rc | Slc30a10 |
| Mtap1b | Gpr133 | Mettl8 |
| Igkv19-93 | Igf1 | Pla2g12a |
| Gm11029 | C1qa | Paqr9 |
| Igkv5-39 | Tle2 | Nqo1 |
| Gm13060 | Leprel2 | 1700006J14Rik |
| Obscn | C1qb | Ninl |
| Ighv1-66 | Arhgef5 | Aqp11 |
| 1300014I06Rik | Pglyrp2 | Gpc4 |
| Gm7592 | BC016579 | Adk |
| Ighg2b | 2010110P09Rik | Casp3 |
| Krt222 | Ctla2b | Reps2 |
| Sspo | Rtp4 | Tfr2 |
| Lrrc32 | Cnrip1 | Tmem20 |
| Igkv3-1 | Sorbs3 | Eda |
| Cd79b | Emr1 | Camsap2 |
| Reln | Unc5a | Hif3a |
| Sdc4 | Tmem121 | 2410127L17Rik |
| Epha2 | Slc22a17 | Hmbs |
| Ccr9 | P2ry14 | Ryk |
| Camk4 | Scn1b | Ppp2r1b |
| Igkv3-5 | Meis1 | Dpf3 |
| Ly6d | Snx24 | Kcng2 |
| Klra17 | 4930539E08Rik | Gna14 |
| Prdm1 | Myct1 | Tgfbr3 |
| Stab1 | Chga | Txnrd2 |
| Ighv7-3 | Ctla2a | Aqp9 |
| Igkj5 | Hoxa5 | Adamts3 |
| Afap1l2 | 8430419K02Rik | Gm15915 |
| Igkv5-43 | Rgs7bp | 4933431E20Rik |
| Sema3a | Pgr | Stard10 |
| Igkc | Emr4 | Blvrb |
| Bach2 | Hmox1 | Ccdc68 |
| Hs3st1 | Pkd2 | 1700001L05Rik |
| Fut1 | Bpifb5 | Selenbp1 |
| Sh2d2a | Kcnd3 | Nefh |
| Fcrl1 | Itih5 | Amigo2 |
| Cd40lg | Oas1g | D16H22S680E |
| Siglech | Gria2 | Fech |
| Rgs4 | Klf12 | Phf10 |
| Cd5 | Gm10021 | Usp15 |

| | | |
|---|---|---|
| Igkv14-126 | Gprc5b | Tspo2 |
| Gm12057 | Naprt1 | Tceal3 |
| Igj | Mn1 | Cpt1c |
| Gm129 | Slc18a1 | 4921530L18Rik |
| Ighv14-3 | 5730409E04Rik | Lpin1 |
| Igkv6-32 | Spic | Slc25a38 |
| Chst2 | Btc | Dnahc12 |
| Cd36 | Slc35d3 | Mc1r |
| Ighv1-4 | Arhgef9 | Armc9 |
| Ighm | Pvrl4 | Hdgf |
| Igkv12-44 | Trim46 | Prelid2 |
| Dlc1 | Zfp608 | Gnb4 |
| Igkv6-15 | Fzd6 | Cd59a |
| Trp53inp2 | Eltd1 | AI427809 |
| Igkv4-61 | Lrch2 | Sdsl |
| Abcc9 | B230378P21Rik | Pitrm1 |
| Cxcr6 | Fn1 | Cacna1g |
| Ppp1r16b | 2210408F21Rik | Bag2 |
| Fcrla | Nap1l3 | Cdr2 |
| Pik3c2b | Hoxa2 | Rasgef1c |
| Adamts5 | Rps6ka6 | 5330416C01Rik |
| Ighv7-1 | Gpx7 | Alad |
| Cd2 | Cpxm1 | Sorbs1 |
| Igkv4-50 | Gm4759 | Narf |
| Itk | Prkag2 | Mpp2 |
| Prickle2 | Gm16197 | Gm10110 |
| Tubb2b | Tanc2 | Zbtb46 |
| Flt4 | Ptpn21 | Slc45a3 |
| Iglv2 | Lpl | Piga |
| Gimap3 | Ccr3 | Col17a1 |
| Il13ra2 | Eya1 | Epor |
| Ets1 | Gm973 | Trim10 |
| Edaradd | Morn4 | Tom1l1 |
| Npnt | Armcx6 | Tjp1 |
| Blk | Mdfi | Lpin2 |
| Igkv10-96 | Odf3b | Prkaa1 |
| Cd3e | Clnk | Syngr1 |
| Sidt1 | Nrxn1 | E2f4 |
| Igkv15-103 | Kitl | Gstm5 |
| Igkv4-91 | Fgf1 | Ext1 |
| S1pr1 | Rbp1 | Gja1 |
| Ighv1-18 | Nhedc2 | Pabpc4 |
| Ighv12-3 | Clmn | Plxnc1 |
| Col5a3 | Tsc22d1 | Uhrf1bp1 |
| Ighv1-26 | Col1a1 | BC057079 |
| Cxadr | E030002O03Rik | Pdlim1 |
| Il2rb | Cd163 | 4632434I11Rik |
| Steap4 | Slc1a3 | Padi3 |
| Cdh5 | Fbn1 | Lonrf2 |
| Ighv3-6 | Gpr171 | Iqcd |
| Cd28 | 2610307P16Rik | Fbxl13 |
| Nid1 | Dmpk | Txnl1 |
| Lrg1 | Tmem44 | Klhl12 |

| | | |
|---|---|---|
| Ppap2b | Glul | Cachd1 |
| Skap1 | Sdc3 | Asb17 |
| Ighv1-62-2 | Gm11428 | Ubxn2b |
| Ighv1-22 | Cd209f | 1700086O06Rik |
| Gimap4 | Pdzk1 | Gm8116 |
| Gpr174 | Cuedc1 | Gm14490 |
| Themis | Dntt | Cela1 |
| Mir5107 | Fgf11 | P2ry1 |
| Acsbg1 | Zfp57 | Alkbh8 |
| Tenc1 | Hdgfrp3 | Urod |
| Amotl1 | Il18 | Aplp2 |
| Ccr6 | Ctxn1 | Cenpv |
| Slc41a2 | Mageh1 | Atp4a |
| Cd38 | Ccdc46 | Uros |
| Sytl2 | Ppp1r26 | Cyp1a1 |
| Bhlhe41 | Mllt4 | Aplf |
| Osmr | Tmem26 | Tarsl2 |
| Epn2 | Ptn | Tbccd1 |
| Rag1 | Myl10 | Btaf1 |
| Tnfrsf25 | Vegfc | Slc17a4 |
| Rasip1 | Akap7 | Ccdc27 |
| Gpr55 | Timd4 | Pvt1 |
| Trem1 | Slc27a2 | Vamp5 |
| Thy1 | Gdf15 | Spry4 |
| Ushbp1 | Btnl4 | Mical3 |
| Galntl2 | Spred3 | Tfrc |
| Tspan7 | Dnalc1 | Ubac1 |
| AC073553.3 | Prrg4 | Cad |
| Il4i1 | Lrp1 | Scin |
| Cd3g | Hoxa6 | Ubxn2a |
| Iglc2 | 6030429G01Rik | Xk |
| Igkv4-57 | Gm10598 | Gdpd1 |
| Tigit | Chl1 | 9430020K01Rik |
| Trbc2 | Flt3 | Vwce |
| Ptprb | Slc25a27 | Wdr55 |
| Traf1 | Frmd4b | Cited4 |
| Grap2 | Tmtc1 | Oat |
| Jag1 | 9930038K12Rik | Wrn |
| Epas1 | Grik5 | Usp24 |
| Trim7 | A730081D07Rik | Il1rl1 |
| Setbp1 | Tgm1 | Fads2 |
| Gpm6a | Ccl4 | Gpsm2 |
| Hspg2 | 1700024P16Rik | 2610034M16Rik |
| Ikzf3 | Myo15 | Nudt9 |
| F8 | 4933409K07Rik | Abca3 |
| Ighv1-53 | Gfra4 | Wdr59 |
| Ighv14-2 | Tmem158 | Daam1 |
| Ighv1-63 | Rhbdl3 | Ahctf1 |
| 2010001M09Rik | Tmem8b | Kif26b |
| Slamf9 | Mycn | Fzd7 |
| Zap70 | C030034L19Rik | Necap2 |
| Kdr | Ctsf | Fam109b |
| Mmrn2 | Tbxas1 | Kctd7 |

Nr2f2
Gbp9
Cd83
Bcar1
Srpk3
Lipg
Kif26a
Ifng
5830411N06Rik
Iglv1
Igkv8-21
Gpr182
Adamts1
Aebp1
C1qtnf1
Chst1
Cd163l1
Gm16603
Pde2a
Ctsl
Nfib
Sparc
Col6a3
Fmo1
Igkv8-16
Igkv5-45
Aldob
Kcnj8
Cldn5
Cd3d
Cyp4b1
Clec2i
Cdr2l
Gm20419
Derl3
Zc3h12d
Parva
Npr1
Grap
Ldb2
Gfra1
Ighv1-49
Rasgrp3
Gm17004
Trbc1
Dgka
Gpr116
Igkv7-33
Siglecg
Lifr
Rasgef1b
Cd97
Pxdn
Lpar1
Smoc1
Ppp2r2b
Kcnj9
Hpgd
Kif5a
Kcnj16
Zan
Nwd1
Gbp11
Slc40a1
Dok7
Rasl12
Nr1h3
Cecr6
Nhsl1
Mrc2
Hba-a2
Cd209g
Vcam1
Hba-a1
Socs5
Havcr2
Alas2
Hbb-b2
Synpo2l
Klhl30
Trim6
Creb5
Zfhx3
6720401G13Rik
Rgag4
Hbb-b1
Rgs1
Gm15893
Gm3892
Mrap
Dcn
Col3a1
Tmem51
Ica1l
Slc39a8
Atp6v0a4
Prdx2
A230051G13Rik
Mtfp1
Leprot
Hebp1
Slc1a4
Paqr4
Cd248
Pkd1l3
Soat2
Pcx
Lyrm4
Slc25a42
Snca
Fn3krp
Sfrp4
BC024659
Rgs9bp
Cyp4f39
Gm11713
March8
Fam158a
Acot6
Cox6b2
Hsph1
Vkorc1l1
Zfp930
Mrpl52
Ccdc23
Dyrk3
Klra8

Atp1b1
Gm15674
Ighv1-59
Galntl4
Ddr1
Sele
Prelp
Dab2
Gm15675
Igkv5-48
Eaf2
Gpr126
H2-DMb2
Shroom2
Igkv2-109
Fermt2
Blnk
Igkv9-120
Tbx21
Pcdh12
Card6
Papln
Card11
Mcam
Fabp4
Gpr18
Tlr1
Sema3f
Adam19
Ctgf
Ighv2-9-1
Hs3st3b1
Clec14a
P2ry10
Cd300c
Cd22
Aplnr
Igkv14-111
Fam46c
Darc
Gramd3
Gpr183
Btg1
Tcf7
Zcchc14
Igfbp7
Cd74
Adcy4
Id3
Cyr61
Iglc3
Ciita
S100a8

Acp5
Prkcb
Nr4a3
Ighv9-2
Igkv4-63
Ighv3-8
Igkv10-94
H2-Eb1
Ighv1-20
Ighv2-2
Ighv14-1
Slamf7
Bmf
Ets2
Hepacam2
H2-Aa
Btla
Chi3l3
Cxcr3
Irg1
Filip1l
Irs2
Cd226
Tlr9
Ccr5
Ighv5-15
March1
H2-Ab1
Bhlhe40
Notch4
Egr3
Dusp10
Fcnb
Ms4a4b
Gm6377
Igkv1-99
